# Cytoplasmic recapping commences the translation of a micropeptide, MARS from MALAT1 after the removal of an inhibitory RNA G-quadruplex

**DOI:** 10.64898/2026.10.13.758904

**Authors:** Avik Mukherjee, Ritesh Sonar, Padmaja Prasad Mishra, Shubhra Majumder, Chandrama Mukherjee

## Abstract

Long non-coding RNAs (lncRNAs), have emerged as key regulators of diverse biological processes, including metabolism, development, cellular differentiation, and stress responses. Most lncRNAs are transcribed by RNA polymerase II and undergo canonical RNA processing, including splicing, polyadenylation, and 5′ N7-methylguanosine (m⁷G) capping and cytoplasmic recapping. However, whether cytoplasmic recapping of lncRNAs can promote the production of functional micropeptides remains poorly understood. In this study, we investigated the functional consequences of cytoplasmic recapping of *MALAT1*, a highly abundant and broadly expressed lncRNA. We demonstrate that cytoplasmic processing and recapping of a *MALAT1* decay intermediate support the production of a previously uncharacterized micropeptide, designated MARS (MALAT1 Responsive to Stress). We further identify RNA G-quadruplex structures in the 5′ region of *MALAT1* that inhibit MARS translation, revealing an additional layer of structural regulation. MARS expression increases in response to arsenite-induced oxidative stress and is associated with nuclear factor erythroid 2-related factor 2 (Nrf2)-containing complexes. Furthermore, MARS is implicated in the regulation of antioxidant response element (ARE) associated genes suggesting a potential role in cellular antioxidant defence. This study highlights a mechanism through which cytoplasmic RNA processing may contribute to micropeptide production and suggests a role for *MALAT1*-derived MARS in oxidative stress-responsive gene regulation.

## Introduction

Every RNA polymerase II (Pol II)-transcribed RNA molecule including all pre-mRNAs, miRNA precursors, long noncoding RNAs (lncRNAs), small nucleolar RNAs (snoRNAs), and small nuclear RNAs (snRNAs) acquires m^7^GpppN cap structure where N is the first transcribed nucleotide at its 5’ end (1). The cap controls every step of RNA metabolism including export to the cytoplasm, translation to mRNA decay. Capping is catalyzed by mRNA Capping Enzyme (CE) bound to the phosphorylated C-terminal domain of Pol II (2). The N-terminal triphosphatase domain of CE generates diphosphate from nascent transcribed triphosphate in the nucleus. Subsequently, the guanyl transferase domain of CE transfers a GMP moiety that is bound to the ε-amino group of lysine 294, to the 5’-end of RNAs, generating a G-capped RNA. RNA guanine-7 methyltransferase (RNMT) in complex with the activator protein adds a methyl group to the N7 position to complete the synthesis of the basic cap structure. This cap structure is sufficient for recognition by cap-binding effector proteins (3) that guide the mRNA to participate in different processes like nuclear processing, export, translation and microRNA silencing, nonsense-mediated mRNA decay and ultimately mRNA turnover (4–6). The long-standing concept of decapping of RNAs led to degradation has been challenged by the discovery of cytoplasmic pool of CE (cCE) in different organisms (7–9).

Although multiple CE homologs encoded by distinct genes in *Trypanosoma* function in nucleus and cytoplasm, mammalian cells possess only one CE-encoding gene (9,10). Mammalian CE exports from the nucleus to the cytoplasm through the Exportin1-mediated pathway (10). Unlike nuclear CE, cCE associates with the cytoplasmic adaptor protein Nck1, and together with an unknown 5’-monophosphate RNA kinase, RNA Methyl transferase (RNMT) and RNMT associated mini activator protein (RAM), to form an active cytoplasmic capping complex (CCC), which restores a cap to uncapped transcripts in the cytoplasm (7,8,11,12). In order to examine the function of cCE, a dominant negative mutant of CE was generated by changing the active lysine residue to alanine (K294A) which was restricted to express only in the cytoplasm (rcCE-K294A or K294A) by introducing a Nuclear Export Sequence (NES) at the N-terminus of CE and deleting the Nuclear Localization Signal (NLS) (7). Overexpressing K294A leads to accumulation of uncapped RNAs and cCE is found to regulate the stability and translability of the target mRNA transcripts by adding cap structures to the uncapped RNAs and thus maintaining ‘Cap homeostasis’ to regulate gene expression (8,9). Advances in sequencing technologies have led to the identification of uncapped RNAs in several organisms (8,9,13). Single cell imaging techniques have further revealed cyclical decapping and recapping of mRNAs suggesting cytoplasmic recapping has broader implications in post transcriptional control (14). Recently, we demonstrated that target mRNAs of cCE regained their cap structures during recovery from brief arsenite-induced oxidative stress (10). Interestingly, cytoplasmic capping has also emerged as an important hub of post-transcriptional regulation for several lncRNAs associated with solid tumors, likely through modulating the cap status of their target transcripts (15). However, the biological significance of cCE in regulating its target lncRNAs and their downstream functions remains unclear. Recently, we showed that cCE contributes in maintaining the stability of several target mRNAs and lncRNAs during CoCl2-induced hypoxia (16,17).

LncRNAs are increasingly recognized as important regulators of cancer development and progression, functioning either as oncogenic drivers or tumor suppressors. lncRNAs share many structural features with mRNAs, including 5’ capping and 3’ poly(A) tails, but generally lack protein-coding capacity. However, accumulating evidence indicates that some lncRNAs can be processed into smaller functional products, and can even generate micropeptides from small ORFs (sORF), typically of 10-100 amino acids. Though the micropeptides were initially unrecognizable and largely overlooked, different research groups across the globe revealed their diverse roles of micropeptides in cellular and developmental processes once the micropeptides were identified though advances in technologies such as ribosome foot printing, mass spectrometry and bioinformatics tools. For example, lncRNA PSMA3-AS1 encodes a 48-amino acid micropeptide, PAMP (proline-associated micropeptide), which acts as a tumor suppressor in lung adenosarcoma (18). Similarly, miPEPDR, another micropeptide derived from lncRNA DANCR, regulates the oxidative stress response in corneal epithelia by modulating PARP1 activity (19). Several antisense lncRNAs or long intergenic RNAs (lincRNAs) have also been shown to encode functional micropeptides. For instance, Linc-mipep and linc-wrb encode micropeptides with homology to the vertebrate-specific chromatin architectural protein Hmgn1, and are implicated in the development of vertebrate-specific brain cell types (20). A recent study has revealed that microproteins that are encoded from sequences with high GC content, which may facilitate non-canonical translation, are susceptible to Cullin-RING E3 ubiquitin ligase (CRL)-dependent proteasomal degradation (21). These findings prompted us to investigate whether cCE regulates the translational potential of its target lncRNAs.

Earlier, we demonstrated that several solid tumours-associated lncRNAs, including *MALAT1, GAS5-838, RP1-28E3.8, RP11-4649.F11,* and *NFYC-AS1* are regulated by cCE (15). In this study, we found that all these transcripts, except *RP11-4649.F11,* may encode micropeptides with diverse subcellular localizations in U2OS osteosarcoma cells suggesting potentially distinct biological functions. Among the cCE-targeted lncRNAs, *MALAT1* was found to contain a functional sORF that is translated exclusively from decay intermediate of the full-length transcript. Notably, a specific RNA G-quadruplex (rG4) motif present at the 5′ end of full-length *MALAT1* transcript appears to inhibit sORF translation. The absence of this motif in the decay intermediate relieves this inhibition, permitting cytoplasmic recapping and facilitating cap-dependent translation of the encoded micropeptide. We named this micropeptide as MARS (MALAT1 Associated Responsive to Stress), since its expression increases during arsenite-induced oxidative stress. Also, our study suggested MARS may act as a transcription regulator in association with nuclear factor erythroid 2-related factor 2 (NEF2L2/NRF2), modulating the expression of stress-responsive genes containing Antioxidant Response Element (ARE) like Superoxide Dismutase 2 (SOD2) and Catalase (CAT).

Collectively, these findings reveal a novel mechanism through which cytoplasmic capping not only stabilizes truncated RNA species but also facilitates the selective translation of micropeptides from lncRNA decay intermediates. This mechanism highlights the expanding role of post-transcriptional RNA modifications in shaping the complexity of the human proteome.

## Materials and Methods

### Data retrieving and bioinformatics analysis

The data of ribosome-bound lncRNAs were retrieved from TransLnc *in silico* tool that provides both computationally predicted and experimentally supported lncRNA peptides (https://bio-bigdata.hrbmu.edu.cn/TransLnc/). Analysis of the Gene Ontology pathway is performed using web servers including DeepGOWeb(22), AmiGO 2(23) and g:Profiler (24).

Three different G4 prediction methods including cGcC (Consecutive G over consecutive C ratio) (25), G4H (G4Hunter)(26) and G4NN (G4 Neural Network)(27) were carried out by G4RNA screener v.0.2 (28) with default setting. The G4 threshold is defined as cGcC >4.5, G4H >0.9, G4NN >0.5.

Anti-oxidant response elements (AREs) within the CAT and SOD2 promoter regions were predicted using the JASPAR database (http://www.jaspar.genereg.net)(29).

### Plasmid constructs and siRNAs

The pEGFP-N1 plasmid (Addgene, USA), a generous gift from Dr. Somshubhra Nath (Presidency University, Kolkata, India), was further engineered for use in this study. The EGFP start codon was mutated to ATT using site-directed mutagenesis strategy of In-Fusion reagent (Takara Bio), generating the pEGFP-N1-mutant vector. To assess the coding potential of candidate sORFs, their sequences were cloned upstream of the mutated EGFP coding sequence in the pEGFP-N1-mutant vector. A corresponding set of constructs was generated in which the start codons of the candidate sORFs were similarly mutated to ATT. Mammalian expression construct of mcherry-H2B, a generous gift from Dr. Benu Brata Das (IACS, Kolkata, India), was used as a co-transfection control. The In-Fusion cloning system (Takara Bio) was used to generate the plasmid constructs described in this study. The 5’-end of full length and decay intermediate of *MALAT1*, the putative rG4 structures located upstream of the predicted *MALAT1* sORFs, and various mutants of these rG4 regions were cloned into the pEGFP-N1-mutant vector. The rG4 sequences were also cloned into the pRL-TK plasmid (Promega) for *in vitro* transcription. The 3×HA-RPL10A construct was cloned into the pcDNA4/TO vector. The HCV IRES was amplified from the pRLH plasmid, a generous gift from Dr. Partha Saha (SINP, Kolkata), and subcloned into the pEGFP-N1-mutant vector. A plasmid expressing GST-eIF4E K119A was obtained from Addgene (USA).

To generate an antibody against MARS, the sORF2 region of MALAT1 was cloned into the pGEX6p-1 vector between the BamH1 and Sal1 restriction sites. The sequences of the primers used in this study are provided in Supplementary Table-1. All plasmid constructs were verified by Sanger sequencing. Silencer select siRNAs targeting MALAT1 (Assay ID-n272231) and XRN1 (Assay ID-s29016), and a Silencer Negative Control siRNA (Assay ID-AM4613) were obtained from Thermo Fisher Scientific, USA.

### Cell culture, treatment, and transfection

U2OS human osteosarcoma cells and human retinal pigment epithelial cells (hTERT-RPE1, referred to as RPE1) were purchased from ATCC, USA, while human epidermal keratinocyte cell line HaCaT was obtained from BRIC-NCCS Cell Repository, India. All cell lines were cultured in Dulbecco’s Modified Eagle Medium (DMEM High Glucose; Gibco, USA) supplemented with 10% Fetal bovine serum (FBS; Himedia, India) and 1% Penicillin-Streptomycin (Gibco). Cells were maintained at 37 °C in a humidified incubator containing 5% CO₂.

JetPrime transfection reagent (PolyPlus) was used to transfect the plasmids into mammalian cells, following the manufacturer’s instruction. siRNAs were transfected at a final concentration of 20 nM using Lipofectamine RNAiMAX (Thermo Fisher Scientific, USA), following manufacturer’s instruction. Briefly, 1.5×10^5^ and 7.5×10^4^ U2OS cells were seeded in 35 mm dish for plasmid and siRNA transfection respectively.

To inhibit cap dependent translation, cells were treated with 20 μM of m^7^G(5’)ppp(5’)G cap analogue (Invitrogen). Oxidative stress was induced by treating cells with 0.5 mM sodium arsenite (NaAsO2, Merck, USA). Cells were collected at the indicated time points during stress and recovery, where the latter was initiated by replacing the arsenite-containing medium with fresh, pre-warmed medium, followed by incubation for an additional 2 h.

### Translating Ribosome Affinity Purification (TRAP)

TRAP assay was carried out according to a previously described method by Reynoso et al (30), with minor modifications. Briefly, U2OS cells expressing HA-RPL10A were resuspended in twice the volume of polysome extraction buffer containing 200 mM Tris–HCl (pH 9.0), 200 mM KCl, 25 mM EGTA (pH 8.0), 35 mM MgCl2, 1% (v/v) Detergent mix, 1% (v/v) PTE, 5 mM DTT, 1 mM PMSF, 50 µg/µL cycloheximide, and 50 µg/µL chloramphenicol). The suspension was incubated on ice for 30 min, and subsequently centrifuged at 16,000 × g for 15 min at 4 °C. A 100 μl aliquot of pre-equilibrated magnetic anti-HA beads (Thermo Fisher Scientific, USA) was added to the cell lysate and incubated for 2 h at 4 °C with gentle back-and-forth rocking or gyration to facilitate binding of the HA-tagged ribosomes to the affinity beads. RNA associated with the bead-bound ribosomes and input fractions were subsequently extracted and qPCR experiments were carried out with gene specific primers. The data was presented as the ribosome bound transcript normalized to the input transcript.

### Whole Cell lysis and isolation of cytoplasmic fraction

U2OS cells were harvested by scraping and pelleted by centrifugation at 1000 × g for 5 minutes. For whole-cell lysate preparation, the cell pellet was resuspended in lysis buffer containing 50 mM Tris-HCl (pH 8.0), 150 mM NaCl, and 1% NP-40, freshly supplemented with 1 mM PMSF and 1X protease inhibitor cocktail (Merck Life Sciences, USA). For subcellular fractionation, the cell pellet was instead resuspended in five volumes of cytoplasmic extraction buffer containing 20 mM Tris-HCl (pH 7.5), 100 mM NaCl, 10 mM MgCl₂, 10 mM KCl, 0.2% NP-40, and freshly supplemented with 1 mM PMSF and 1× protease inhibitor cocktail. In both procedures, the resuspended cells were kept on ice for 10 minutes with gentle agitation by occasional gentle tube flicking to facilitate complete lysis.

For subcellular fractionation, the lysates were centrifuged at 1000 × g for 10 minutes at 4°C. The nuclear fraction was recovered in the pellet and the supernatant was carefully transferred to a fresh tube, which was used as the cytoplasmic fraction. For whole-cell lysate preparation, the lysate was centrifuged at 10,000 × g for 10 minutes at 4°C to remove the cellular debris. The total protein concentration in each lysate was determined using the BCA protein assay kit (Thermo Fisher Scientific).

### Cytoplasmic RNA extraction

Cytoplasmic RNA was extracted from the cytoplasmic fractions using Trizol reagent (Invitrogen, USA) following the manufacturer’s protocol. RNAs were treated with DNase I (Thermo Fisher Scientific, USA), and recovered using phenol:chloroform:isoamyl alcohol (SRL) extraction following a published protocol (31).

### Recombinant protein expression, purification and antibody generation

GST-eIF4E K119A was expressed in *Escherichia coli* BL21 (DE3), induced by 0.5 mM isopropyl-β-d-thiogalatopyranoside (IPTG) that was added when OD600 of the bacterial culture reached 0.6, and subsequently incubated overnight at 16 °C. The harvested cells were lysed by resuspending in GST lysis buffer containing 20 mM Tris-Cl (pH 8), 20 mM NaCl, 1 mM DTT, 1 mM PMSF, 1% Triton X-100, 125 μg/ ml lysozyme and 5% glycerol, and sonication. After centrifugation at 15,000 × g for 30 minutes at 4°C to remove the debris, the supernatant was applied to glutathione Sepharose 4B beads (GE Healthcare) for affinity purification. Following extensive washing with GST wash buffer containing 25 mM Tris-Cl (pH 7.5), 150 mM NaCl and 0.1% NP40, the bead-bound GST-eIF4E K119A was stored in bead storage buffer (20% glycerol and 1 mM PMSF in GST-wash buffer) until further use. The recombinant plasmid expressing GST-sORF2 was given to Biobharati Life Sciences Private Limited, India, for the generation and purification of rabbit polyclonal antisera against MARS.

### Separation of capped and uncapped transcripts with GST-eIF4E protein

Capped and uncapped RNAs were separated using a method described previously (12). Briefly, 4 μg of bead bound GST-eIF4E K119A was equilibrated in Cap PD buffer containing 10 mM Tris (pH 7.5), 150 mM NaCl, 0.1% (v/v) IGEPAL CA-630, and 1 mM DTT. Test RNA samples were prepared with 1 μg cytoplasmic RNA, 0.04 ng each of capped luciferase and uncapped GFP RNA as spiked in controls along with 80 U RNaseOUT (Thermo Fisher) in Cap PD Buffer. Equilibrated GST-eIF4E K119A protein-bound beads were added to RNA samples and incubated end-over-end at 4°for 2 h. The flow through was collected to isolate the uncapped RNA pool by Trizol and the beads were washed thrice with 200 μl Cap PD Buffer at 4°C for 5 min each. Washed beads were resuspended in 200 μl Trizol and purified RNA from the flow through or bound fractions were eluted in 15 μl of nuclease free water.

### *In vitro* transcription

Candidate RNA G4 oligos were generated by *in vitro* transcription MEGAShortScript T7 kit (Invitrogen, Thermo Fisher Scientific, USA), following the manufacturer’s instructions. Briefly, RNA segments cloned into the pRL-TK vector were linearized using NotI restriction enzymes. 1 μg of purified linearized plasmid DNA was used as a template for in vitro transcription. *In vitro* transcribed RNAs were purified using the Monarch Spin RNA Cleanup kit (New England Biolab, USA).

### Circular dichroism (CD) spectroscopy

CD assay was performed with a Chirascan Plus CD spectrometer of detection range of 150-850 nm (Applied Photophysics, UK) equipped with a 1 cm path length quartz cuvette. The reaction samples containing 5 μM oligos were prepared in 10 mM Tris-Cl buffer (pH 7.0) and 150 mM KCl/LiCl to form a total volume of 2 ml. The mixtures were then vortexed and heated at 95°C for 5 min and allowed to cool for 15 minutes at room temperature for renaturation. The samples were scanned at a 2 nm interval from 220 to 310 nm, and the data were blanked and normalized to mean residue ellipticity before smoothed over 10 nm. The raw data generated is in the KIN file format. All the data were taken from the instrument for processing through Origin 2018 and were further analyzed with Spectra Manager™ Suite and Microsoft Excel.

### Thioflavin T (ThT) Fluorescence Assay

The Thioflavin T (ThT) fluorescence assay was performed as described previously (32). Briefly, candidate oligonucleotides were prepared at a final concentration of 4 μM in 20 mM Tris-HCl buffer (pH 7.0) containing 40 mM KCl, with a final reaction volume of 100 μL. The samples were vortexed followed by heating at 95°C for 5 min, and allowed to cool to room temperature for 15 min to allow the oligonucleotides to refold. ThT was then added to a final concentration of 2 μM, and the samples were mixed thoroughly. Fluorescence emission was recorded over a range of 450-700 nm at 50 nm intervals using a multimode plate reader (Synergy, Agilent technologies, USA). The fluorescence profiles were compared to assess ThT binding and the potential of the oligonucleotides to adopt rG4 structures.

### Immunoprecipitation

Whole cell extracts from U2OS cells were used for immunoprecipitation with magnetic Protein G beads (Bio-Rad, USA). Briefly, 40 µl bead suspension was equilibrated in IP buffer containing 25 mM Tris-HCl, (pH 7.5), 150 mM NaCl, and then blocked with blocking buffer consisting of 1% BSA in IP buffer, for 30 min. The beads were then incubated with 4 μg mouse anti-GFP antibody (Abclonal, USA) for 1 h at room temperature with shaking. Cell extracts were pre-cleared by incubation with 40 μL of beads for 2 h at 4 °C. Next, the antibody-bound beads were added to the pre-cleared cell extracts and incubated overnight at 4°C with shaking. The beads used for pre-clearing the extracts were used as the negative control. Both the control and antibody-bound beads were extensively washed with IP wash buffer containing 0.1% NP40 in IP buffer. The bead-bound proteins (100%) and 5% of the input samples were analyzed by immunoblotting.

### Western blotting

Protein samples were analyzed by SDS-PAGE, and transferred onto Immobilon-FL PVDF membrane (Millipore, USA). Membranes were blocked with 3% BSA (SRL, India) in phosphate-buffered saline (PBS) for 30 min at room temperature, and incubated overnight at 4°C with primary antibodies diluted in the blocking buffer. The primary antibodies used in this study were rabbit anti-GFP (1:10000, #A11122, Invitrogen, USA), mouse anti-mCherry (1:5000, #68088-1-Ig, Proteintech, USA), rabbit anti-Xrn1 (1:1000, #PA5-57110; Invitrogen, USA), mouse anti-α-tubulin (1:5000 dilution, #SC-17787; SCBT, USA), rabbit anti-NRF2 (1:1000, #A0674, Abclonal, USA), rabbit anti-HA (1:1000, #3724 CST, USA), mouse anti-human-β actin (1:5000, #AC004, Abclonal, USA). After incubation with primary antibodies and subsequent washing in PBS with 0.1% Tween-20 (PBS-T), the membranes were kept in secondary antibodies DyLight 800-conjugated donkey anti-mouse-IgG (Invitrogen, USA), or Alexa Fluor 680-conjugated donkey anti-rabbit-IgG (Invitrogen, USA) at 1:10,000 dilution in PBS-T. The immunoblots were scanned using an Odyssey CLx infrared imaging system (Li-cor, USA), and the background-corrected intensities of bands were determined using ImageStudio (Li-cor, USA).

### Indirect immunofluorescence

Cells were grown on coverslips and fixed for 10 min at room temperature in a buffer containing 4% paraformaldehyde (Electron Microscopy Sciences, USA) and 0.2% Triton X-100 (Bio-Rad, USA) in 1× PBS. After washing with 1× PBS containing 0.5 mM MgCl2 and 0.05% Triton X-100, cells were incubated in a blocking buffer containing 5% FBS, 0.2 M glycine, 0.1% Triton X-100 in 1× PBS) at room temperature for 30 min. Subsequently, the cells were incubated overnight at 4°C with the primary antibodies diluted in the blocking buffer. The cells were incubated with the primary antibody (rabbit anti-GFP, #A11122 Invitrogen, USA (1:2500); rabbit anti-NRF2, #A0674 Abclonal, USA (1:200); mouse anti-c-myc, #32293 SCBT, USA (1:200); goat anti-eIF3b, #SC-16377 SCBT, USA (1:1000); rabbit anti-MARS, Bio Bharati LifeSciences India (1:200) for overnight at 4°C. After washing, the cells were incubated with donkey anti-rabbit IgG, or donkey anti-mouse-IgG secondary antibodies conjugated to either Alexa Fluor 488 or Alexa Fluor 568 (1:1000; Invitrogen, USA), and Hoechst 33358 (Sigma, USA) diluted in the blocking buffer, for 45-60 min at room temperature. FM4-64FX (1µg/µl, F34563, Thermo-Fischer, USA) was used to stain the cellular membrane. The coverslips were mounted on glass slides using SlowFade mounting medium (Invitrogen, USA). Cells were washed extensively using wash buffer, and then were incubated at room temperature for 60 min with secondary antibodies that were donkey anti-rabbit tagged with either Alexaflour488 or Alexafluor568 (1:1000; Thermo Fisher Scientific, USA), and Hoechst 33358 (Merck Life Sciences, USA) diluted in the blocking buffer. The coverslips were mounted on glass slides using SlowFade mounting medium (Invitrogen, USA). Images were acquired using a confocal laser scanning microscope (Leica, Germany) fitted with a 63× Plan Apo oil immersion objective (NA 1.4). Images were analyzed, and the fluorescence intensity measurements were performed using associated Las-X software package.

### Reverse Transcription-Quantitative PCR (RT-qPCR)

Poly(A)-selected RNAs reverse transcribed using oligo(dT) primer and Super Script III First Strand cDNA Synthesis kit (Thermo Fisher Scientific, USA). Primers targeting the 5′ ends of the selected transcripts and spike-in controls were designed, which are provided in the Supplementary Table-1. qPCR was performed with SsoAdvanced Universal SYBR® Green Supermix (Bio-Rad, USA) using a CFX Connect real-time PCR instrument (Bio-Rad, USA). The absolute quantification of the target genes was performed according to the standard protocol described in Bio-Rad Real-Time PCR application guide. Graphs were generated using GraphPad Prism V5 and statistical significance was assessed using an unpaired, two-tailed Student’s t-test.

### Absolute quantification of *MALAT1* transcript variants

Primer pairs were designed to distinguish full-length *MALAT1* transcript from the decay intermediate. One primer pair targeted the 5′ region of *MALAT1*, which is present in the full-length transcript but absent from the decay intermediate. A second primer pair targeted a region common to both transcripts. Ct values obtained using the two primer pairs were used to determine the absolute copy numbers based on the standard curve method described in Bio-Rad Real-Time PCR application guide. The copy number obtained using the common-region primers represented the total *MALAT1* transcript population, whereas the 5′-specific primers measured full-length *MALAT1* transcript. The copy number of the decay intermediate was calculated by subtracting the copy number of full-length *MALAT1* from the copy number of total *MALAT1*.

### Molecular docking

The 3D structure of Nrf2 was retrieved, and the 3D structure of MARS was predicted, constructed and downloaded from AlphaFold 3 database. The structure of the Antioxidant Response Element (ARE) and Hypoxia Responsive Element (HRE) were constructed from their respective consensus sequences and built using UCSF ChimeraX software. Full length NRF2 and MARS, NRF2 and ARE, NRF2 and HRE were subjected to molecular docking using HADDOCK 2.4. The predicted interaction surfaces were analyzed using PDBePISA. PyMOL software was used to visualize, modify and export the PDB files.

### Chromatin immunoprecipitation (ChIP) assay

Chromatin immunoprecipitation (ChIP) was carried out following the protocol described by Katsouda et al (33). Briefly, cells expressing either EGFP or MARS-EGFP were cross-linked with 1% formaldehyde for 10 minutes at room temperature to stabilize DNA–protein interactions, and the reaction was quenched with glycine. Cells were then washed with ice-cold PBS containing 1 mM PMSF and harvested by centrifugation at 2000 × g for 5 mins at 4 °C. The cell pellet was lysed in lysis buffer (1× buffer A, 0.5 mM DTT, 1× protease inhibitor cocktail, 1 mM PMSF), and nuclei were collected by centrifugation at 2000 × g for 5 minutes at 4 °C. The buffer A consists of 1% SDS, 10 mM EDTA (pH 8.0), 50 mM Tris (pH 8.1) and 0.1% Triton X-100. The nuclear pellet was resuspended in buffer B [50 mM Tris-HCl (pH 8.1), 100 mM NaCl and 5 mM CaCl₂] supplemented with 0.5 mM DTT and subjected to micrococcal nuclease digestion for 20 mins at 37 °C. Following centrifugation at 16,000 × g for 1 min at 4 °C, the nuclei were resuspended in ChIP buffer (1× ChIP buffer, 1× protease inhibitor cocktail, 1 mM PMSF) and sonicated to disrupt nuclear membranes. The ChIP buffer is composed of 0.1% SDS, 1 mM EDTA (pH 8.0) and 10 mM Tris-HCl (pH 8.1). The lysate was clarified by centrifugation at 9400 × g for 10 mins at 4 °C. Approximately 10 µg of chromatin was incubated overnight at 4 °C with either rabbit anti-GFP antibody (Bio Bharati Life Sciences, India, cat# 36169) or normal rabbit IgG (Bio Bharati Life Sciences, India, cat# BB-AB003) as a negative control. Protein G magnetic beads were then added to the immunoprecipitation samples and incubated for 2 hrs at 4 °C with rotation. The bead– chromatin–antibody complexes were sequentially washed three times with low-salt wash buffer and once with high-salt buffer, followed by elution at 65 °C for 30 mins. Proteins were digested using Proteinase K at 65 °C for 2 hrs, and DNA was purified using a spin column. Both immunoprecipitated and input DNA samples were analyzed by agarose gel electrophoresis and quantitative real-time PCR using SsoAdvanced Universal SYBR Green Supermix (Bio-Rad, USA cat# 172–5271) on a CFX Connect Real-Time PCR system (Biorad, USA). Relative fold enrichment was calculated using the comparative Ct (ΔΔCt) method, with rabbit IgG serving as the negative control.

### Statistical analysis

Statistical analyses were performed using GraphPad Prism 9 (GraphPad Software, Inc.). Differences between two groups were assessed using an unpaired, two-tailed Student’s t-test, while comparisons among multiple groups were performed using one-way ANOVA followed by Tukey’s multiple comparisons test. Data are presented as mean ± SD from at least three independent biological replicates (n ≥ 3). A p-value ≤ 0.05 was considered statistically significant, and n.s. indicates p> 0.05 (non-significant).

## Results

### cCE targeted lncRNAs contain translatable sORFs

Our earlier study identified several lncRNAs regulated by cytoplasmic capping. Notably, the uncapped forms of these lncRNAs were found to be stable. Given the increasing evidence that lncRNAs can encode micropeptides from small ORFs (sORFs), we wondered whether these cCE-regulated lncRNAs can encode micropeptides as well.

To identify cCE-regulated lncRNAs with potential protein-coding capacity, we analyzed publicly available ribosome profiling (Ribo-Seq) datasets (34). This analysis revealed ribosome occupancy on several previously identified cCE-targeted lncRNAs including *MALAT1, GAS5, RP1 and NFYC-AS1*, suggesting their potential association with the translational machinery. In contrast, another cCE targeted transcript, *RP11-464F9.11*, showed markedly low ribosome occupancy (Fig. 1A). Furthermore, TransLnc analysis (34) identified multiple predicted sORFs distributed across the gene bodies of these lncRNAs (Fig. 1B). For simplicity, these sORFs were written as ORFs throughout the text and the figures.

**Figure 1.**
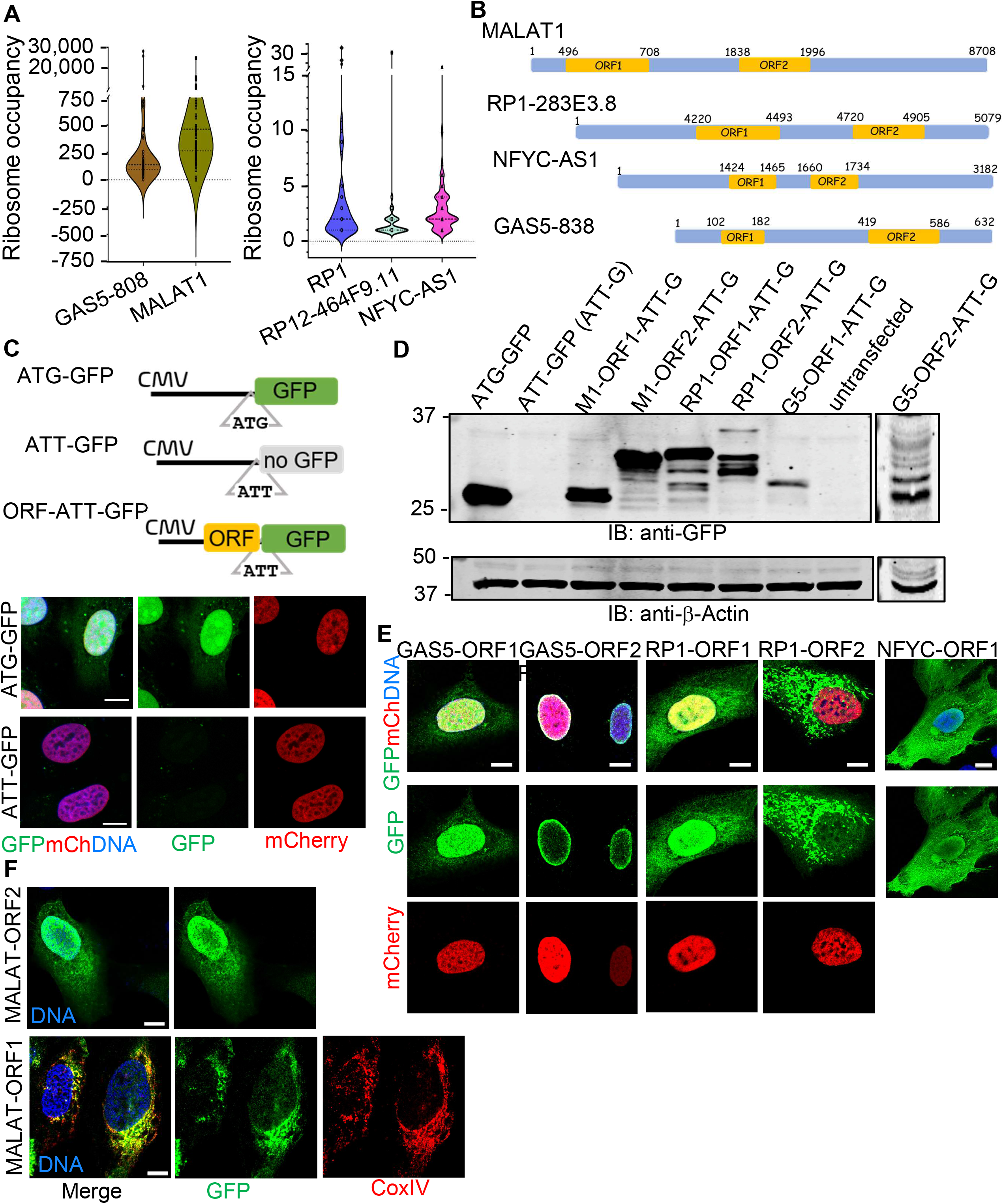
Identification of cCE targeted lncRNA encoded small ORF encoded micropeptides. (A) Distribution of the ribosome occupancy of candidate cCE transcripts as obtained after analysis through Translnc tools, shown as violin plots. The central lines indicate the median and the internal marks represent the distribution of individual observations. (B) Schematic representation of the cCE targeted lncRNAs indicating the relative positions of their sORFs. For simplicity, sORFs are denoted as ORFs in the figures. (C) Schematic of the reporter constructs used to assess sORF-encoded micropeptide translation. ATG-GFP represents the wild type EGFP construct, ATT-GFP represents the mutated EGFP and an ORF-EGFP fusion reporter in which the candidate ORF is positioned upstream of mutant EGFP. For simplicity, EGFP is denoted as GFP in the figures. mCherry-Histone 2B (mCh-H2b) is used as a transfection control. DNA was stained to visualize nuclei. (D) Immunoblot analysis of ORF-GFP chimeras expressed from the indicated reporter constructs. β-Actin is served as the loading control. Molecular weights of the proteins are indicated on the left. (E) Representative micrographs showing GFP and mCherry-H2B signals in cells expressing the chimeric GFP constructs for each of the candidate ORFs. DNA was stained to visualize nuclei. (F) Representative micrographs demonstrating the intracellular distribution of the MALAT1 ORF1 and ORF2 chimeras. Mitochondria and nuclear DNA are stained by CoxIV and Hoechst 33358 respectively. Scale bar is 10 μm.

To experimentally evaluate the coding potential of the predicted ORFs, we used pEGFP-N1 plasmid, where EGFP (a brighter GFP) expression is initiated from its native start codon (ATG-GFP). In contrast, the engineered pEGFP-N1-mutant construct, where the EGFP start codon was changed to ATT (ATT-GFP), does not express EGFP (Fig. 1C). To test the translatability of candidate ORFs derived from *MALAT1, GAS5, RP1* and *NFYC-AS1*, the predicted ORFS were cloned upstream of ATT-GFP, resulting in chimeric ORF-ATT-EGFP constructs, except ORF2 of *NFYC-AS1* (Fig. 1C). Consequently, the expression of EGFP from the mutant constructs would indicate that the upstream candidate ORF can initiate translation to produce a fusion protein containing EGFP (Fig. 1C). The resulting constructs were transfected into U2OS cells and fusion protein expressions were evaluated using immunoblotting and immunofluorescence microscopy (IF). All tested ORF-ATT-GFP constructs produced fusion proteins, with expression comparable to that of the wild type GFP, supporting the translatability of the tested ORFs (Fig. 1D).

We next investigated the subcellular localization of the candidate ORF-derived fusion proteins in U2OS cells. The constructs were co-transfected with a plasmid expressing mCherry-H2B, which served as a nuclear marker. The fusion proteins exhibited distinct subcellular localization patterns (Fig. 1E-F). While GAS5-ORF1 was predominantly nuclear, GAS5-ORF2 showed a perinuclear localization. Similarly, RP1-ORF1 was mostly nuclear, whereas RP1-ORF2 was predominantly cytoplasmic (Fig. 1E). MALAT1-ORF1 predominantly localizes at the mitochondria, as indicated by colocalization with CoxIV, whereas MALAT1-ORF2 displayed a different localization, which is distributed between the nucleus and cytoplasm (Fig. 1F). Notably, NFYC-AS1-ORF1 exhibited a localization pattern distinct from those of the other GFP chimeras (Fig. 1E). Such diverse subcellular distributions of different ORF-derived proteins from the same or different lncRNA suggest that these candidate micropeptides may have distinct cellular functions.

Among the identified cCE-targeted lncRNAs, *MALAT1* exhibited substantial ribosome footprint density (Fig.1B). We therefore selected *MALAT1* as a model transcript to investigate the mechanisms regulating the expression of lncRNA-derived ORFs and to determine how cytoplasmic capping contributes to the generation of the micropeptides. The ORFs predicted by Translnc tool utilizes OpenProt pipeline, which employs ATG as initiation codon (35). To further verify whether the expression of the *MALAT1* ORFs-derived micropeptides depended on the initiation codon, we generated new constructs where the ATG start codons of ORF1 and ORF2 were mutated to ATT (Fig. S1A). Expression of the wild type and mutated fusion constructs of MALAT1-ORFs chimeras were subsequently examined using both immunoblotting and IF studies (Fig. S1). Mutation of the initiation codons resulted in the loss of expressions of both MALAT1-ORF1 and MALAT1-ORF2 constructs (Fig. S1B-C), confirming the need for ATG as the translation initiation codon for both ORFs.

### MALAT1-ORF2 is expressed from the decay intermediate of parental transcript

During our investigation of lncRNAs as substrates of cCE, we identified uncapped forms of several stable decay intermediates of *MALAT1* in U2OS cells, which were subsequently cloned and sequenced (15). Among these, the largest decay intermediate, lacking the 5’ end of *MALAT1* transcript, was identified by cloning and sequencing of the 5’ rapid amplification of cDNA ends (5’ RACE) products (15) (Fig. S2). Interestingly, analysis using the Translnc tool revealed strong ribosome association of this decay intermediate (Fig. 1A). Notably, a major portion of MALAT1-ORF2 (nucleotides 1838-1935), which encodes a 52 amino acids peptide that we named MARS, lies within this decay intermediate (Fig. 2B, Fig. S2). These observations prompted us to investigate whether transcript architecture influences the expression of MALAT1-ORF2-derived micropeptide.

**Figure 2.**
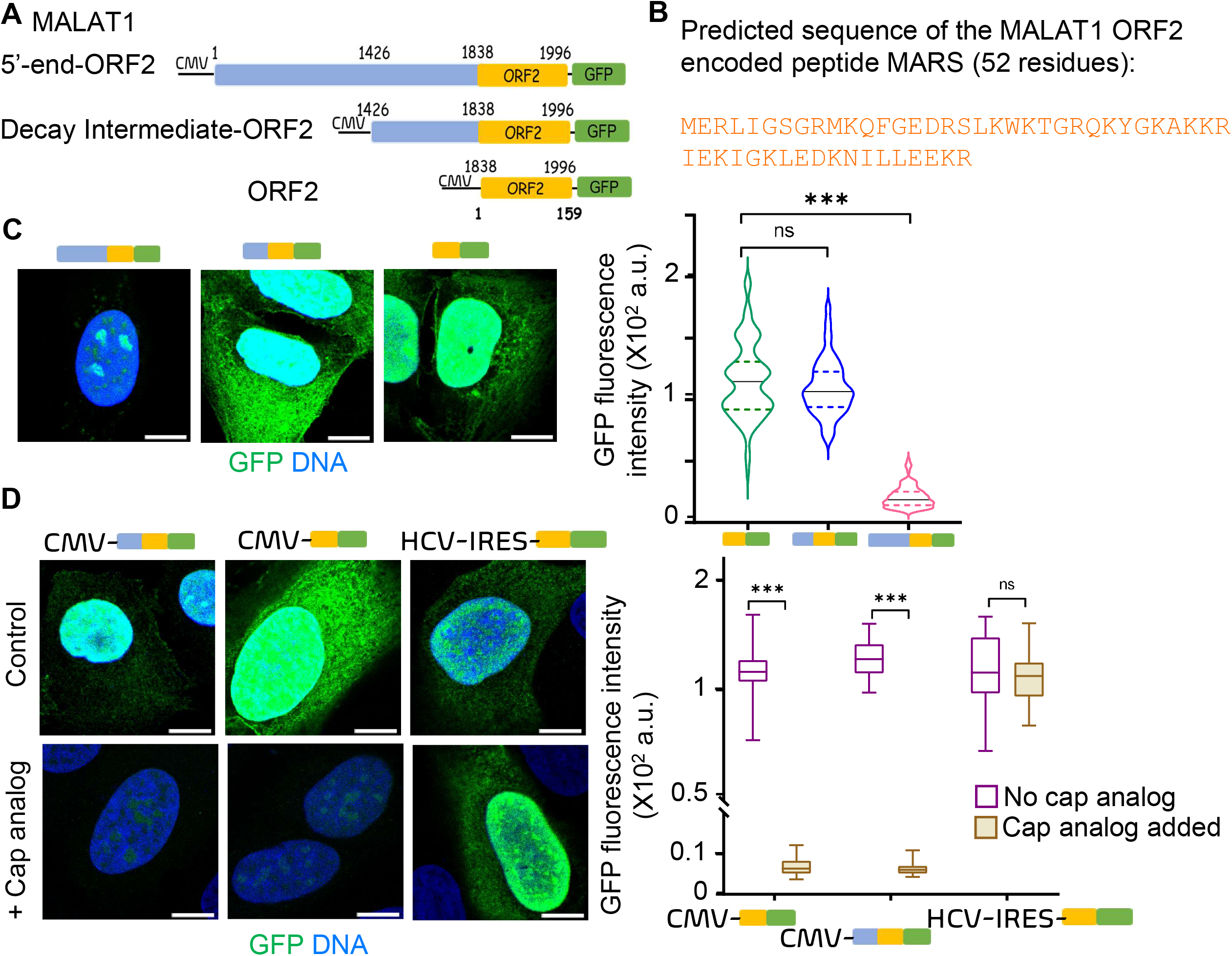
MARS, the micropeptide encoded by of MALAT1 ORF2 is generated via Cap dependent translation. (A) Various constructs of MALAT1-ORF2 with the 5′-end and the decay-intermediate or lacking both are cloned upstream ATT-GFP reporter construct. (B) Predicted amino acid sequences of the MALAT1 ORF2-encoded peptide, designated as MARS, comprising 52 amino acid residues. (C) Representative micrographs show GFP fluorescence from cells expressing either from ORF2-GFP or Decay intermediate-GFP chimeras. No signal is observed from MALAT1 -ORF2 with 5’ end. Violin plots show the quantification of GFP fluorescence intensity (×10² arbitrary units) from the transfected cells (n=30) with the indicated reporter constructs. (D) Schematics illustrate the Decay intermediate-ORF2 and ORF2 constructs cloned under the CMV promoter, as well as the ORF2 construct containing an upstream HCV-IRES in the ATT-GFP reporter construct. Cells are transfected with the above constructs in the presence or absence of cap analog. Representative micrographs showing GFP fluorescence from the CMV driven promoters in absence of cap analog (Control) and reduced expression in presence of cap analog. Expression of GFP driven from HCV-IRES does not alter in presence or absence of cap analog. Scale bar is 10 μm. DNA is in blue. GFP fluorescence intensity is measured from atleast 30 cells and plotted. Statistical significance is indicated by asterisks (*P < 0.05; **P < 0.01; ***P < 0.001) and ns represents non-significant.

To understand this in the context of the native transcript, sequences corresponding to the full length *MALAT1* transcript (1–1996) and the decay intermediate (1426–1996) containing ORF2 were independently cloned upstream of the ATT-GFP cassette in the pEGFP-N1-mutant vector (Fig. 2A). These constructs were designed to assess the influence of upstream *MALAT1* transcript sequences on ORF2 translation. Surprisingly, IF analysis revealed a marked reduction in ORF2-derived EGFP reporter expression from the construct containing the full-length *MALAT1*, compared to robust reporter expression from the construct containing the decay intermediate, which was comparable to that observed with the ORF2-GFP construct alone (Fig. 2C).

We next investigated whether translation of the MALAT1-ORF2 was dependent on the presence of a 5′ cap. Accordingly, U2OS cells expressing the decay intermediate-ORF2-ATT-GFP and ORF2-ATT-GFP constructs were treated with an m7G cap analogue. This analogue competitively inhibits the interaction between the cap binding protein eIF4E and the 5’ m7G cap of transcripts, thereby interfering with global cap-dependent translation. Cap analogue treatment significantly reduced the EGFP expression from both the decay intermediate-ORF2-ATT-GFP and ORF2-ATT-GFP compared with the corresponding untreated controls (Fig. 2D). These results suggest that robust ORF2 translation is indeed cap dependent. To further assess the specificity of this effect, we cloned the hepatitis C virus derived Internal Ribosomal Entry Site (HCV-IRES) upstream of ORF2-ATT-GFP reporter and monitored EGFP expression in presence or absence of cap analogue. Expectedly, cap analogue treatment did not significantly attenuate EGFP expression from the IRES-containing construct, consistent with the ability of the IRES to mediate translation initiation independent of 5’ cap (Fig. 2D).

Overall, these findings suggest that sequences present at the 5’ end of the full-length *MALAT1* transcript repress sORF2 translation, and that processing of *MALAT1* into a stable decay intermediate may facilitate the cap-dependent translation of the sORF2, expressing micropeptide MARS. This observation reveals a novel post-transcriptional regulatory mechanism, through which an embedded ORF of *MALAT1* transcript produces a micropeptide production from *MALAT1*.

### rG4 motif located upstream of *MALAT1* ORF2 inhibits its translation

To gain insight into the mechanism underlying the inhibition of ORF2 translation from the full-length *MALAT1* transcript, we investigated whether RNA secondary structures located upstream of the ORF2 contribute to this. Previous studies have identified a unique bipartite triple-helix structure and several rG4 motifs within *MALAT1*, predominantly in its 3′ region, where they can interact with regulatory proteins and modulate the stability and function of *MALAT1* (34–36). We therefore sought to determine whether additional rG4-forming sequences are present upstream of ORF2.

Computational analysis using the *in silico* G4-RNA Screener tool identified two candidate sequences, designated MALAT1_1 and MALAT1_2, upstream of ORF2, with a high propensity to adopt rG4 conformations containing bulges (Fig. 3A). Based on their position within *MALAT1* relative to ORF2, these candidate motifs were designated the distal and proximal rG4s, respectively (Fig. 3A). To experimentally validate their ability to form rG4 structures, circular dichroism (CD) spectroscopy using *in vitro* transcribed RNA oligonucleotides corresponding to the candidate sequences were performed. A RNA oligonucleotide containing a well-characterized rG4-forming sequence from *NRAS* (37), was used as a positive control, whereas a non rG4-forming sequence of *MALAT1* (1207–1241) served as a negative control. Under physiologically relevant potassium ion concentrations (150 mM K^+^), both the proximal and distal RNA oligonucleotides exhibited characteristic CD spectra, with a negative peak (trough) at approximately 240 nm and a positive peak (crest) at approximately 260 nm. These spectral features were similar to those observed for the *NRAS*-rG4, indicative of a parallel-stranded rG4 topology. In contrast, the non rG4 forming oligonucleotide exhibited a completely different spectral profile (Fig. 3B). Notably, the observed CD signatures were dependent on the presence of K^+^ ion. Replacing K^+^ with Li^+^ (150 mM), which is less favourable for G-quadruplex formation (38), significantly disrupted the characteristic spectral features (Fig. 3B). However, the loss of the spectral features observed for the *MALAT1* distal rG4 was less pronounced than that of the *MALAT1* proximal rG4 in presence of LiCl, consistent with previous observation that not all rG4 structures are equally perturbed by Li^+^ ions (39).

**Figure 3.**
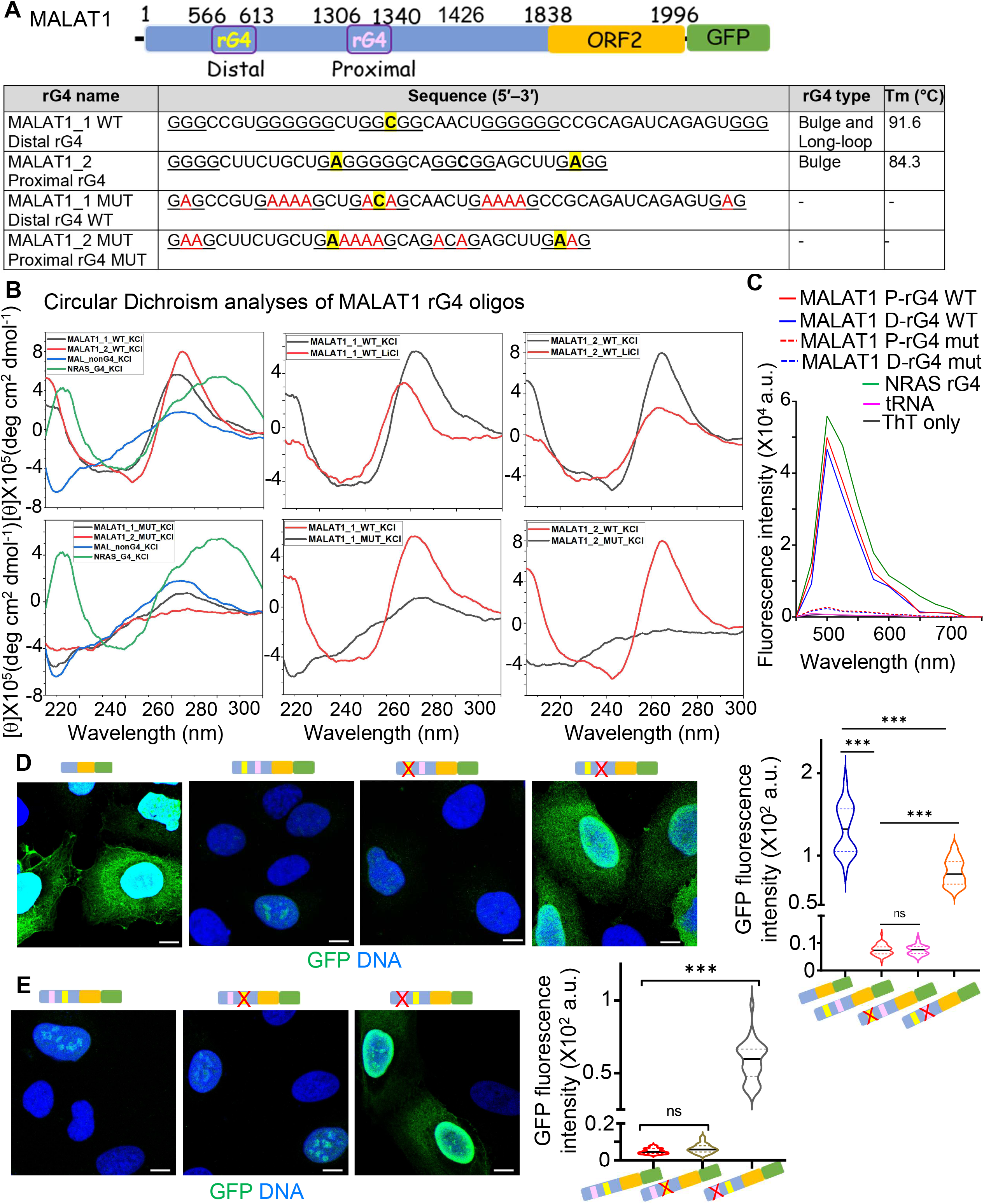
rG4 structure at the 5′ end of *MALAT1* upstream of ORF2 impede its translation. (A) Schematic representation of the *MALAT1* region containing two predicted rG4 structures, located upstream to the ORF2 and termed distal rG4 and proximal rG4, based on their distance from ORF2. The sequences of the wild-type and mutant oligonucleotides used in the study are in the table below. The highlighted residues indicate the bulge. The mutations (indicated by red) were introduced into the G-rich regions to disrupt the corresponding rG4 structures. The predicted rG4 type and melting temperature (Tm) for the wild-type oligonucleotides are indicated. (B) Circular dichroism (CD) spectra of the *MALAT1* rG4 oligonucleotides and their corresponding mutant sequences. Spectra were recorded over the indicated wavelength range to examine the secondary structures formed by the wild-type and mutant oligonucleotides. *NRAS* rG4 was included as a reference for rG4 sequence. (C) Thioflavin T (ThT) fluorescence analysis of the *MALAT1* rG4 wild type and mutated oligonucleotides obtained after incubation with ThT. *NRAS* rG4 was used as a positive control, while tRNA and ThT alone were included as controls. (D) Representative micrographs showing GFP expression from the reporter constructs containing the indicated *MALAT1* rG4 sequences. The corresponding GFP fluorescence intensity of the reporter constructs is quantified on the right. (E) Representative micrographs showing expression of GFP from cells transfected with swapped rG4 reporter constructs and their mutants. Quantification of GFP expression from reporter constructs carrying the indicated rG4 mutations (cells analysed≥30) are plotted as violin plots. For both D and E, nuclear DNA shown in blue. Statistical significance is indicated by asterisks *P < 0.05; **P < 0.01; ***P < 0.001 and ns represents non-significant. The scale bar corresponds to 10 μm.

To further confirm rG4 formation by the distal and proximal motifs of *MALAT1*, we incubated the corresponding RNA oligonucleotides with Thioflavin T (ThT), a fluorescent probe commonly used to detect G-quadruplex structures due to its specific binding to rG4 structures (32). ThT fluorescence was high in presence of control *NRAS rG4,* whereas the presence of *E.coli* tRNA did not show any ThT fluorescence, supporting the preferential recognition of rG4 structures by ThT under the conditions tested (Fig. 3C). In this condition, both distal and proximal rG4 motifs of *MALAT1* showed high ThT fluorescence, suggesting that both these motifs can adopt rG4 structures.

To further establish the structural specificity of the identified rG4 motifs, we generated mutant variants of both sequences by introducing nucleotide substitutions predicted to disrupt G-quadruplex formation while minimizing changes to the overall sequence context (Fig. 3A). CD spectroscopy revealed that, unlike their wild-type counterparts, the mutant proximal and distal rG4s lacked the characteristic CD signature of parallel rG4 structures under physiologically relevant potassium ion concentration, indicating impaired G-quadruplex folding (Fig. 3B). Also, there was a substantial drop in ThT fluorescence signal in presence of these mutant motifs (Fig. 3C), confirming that both upstream sequences upstream of ORF2 can form stable parallel-stranded rG4 structures *in vitro*. We next investigated the functional contribution of these two rG4 motifs to ORF2 translation. To this end, we generated mutant versions of the previously described full-length MALAT1-ATT-GFP reporter constructs, where each rG4 motif was separately disrupted. The constructs, along with the wild type one, were transiently expressed in U2OS cells, and ORF2-derived EGFP expression was subsequently assessed. Disruption of the proximal rG4 resulted in a partial restoration of ORF2 expression relative to the decay intermediate-ORF2 construct, suggesting that the proximal rG4 structural exerts a repressive effect on ORF2 translation (Fig. 3D). In contrast, mutation of the distal rG4 had no detectable effect on reporter expression, suggesting that this motif does not substantially contribute to the regulation of ORF2 translation (Fig. 3D).

To determine whether the observed regulatory effect depended on the position of the rG4 within the transcript or on the sequence-specific intrinsic properties of the motifs itself, the positions of the proximal and distal rG4 motifs were exchanged. This approach generated constructs in which the proximal rG4 was relocated to the original position of the distal rG4 and vice versa (Fig. 3E). We then selectively disrupted each rG4 motif in the position-swapped constructs, and expressed the resulting constructs in U2OS cells. However, the position-swapped construct did not express the ORF2-derived reporter EGFP (Fig. 3E, the first panel). Notably, disruption of the relocated proximal rG4 led to partial restoration of ORF2 expression despite its increased distance from the ORF2 region (Fig. 3E, the last panel). However, disruption of the distal rG4 did not affect ORF2 expression, even when the motif was repositioned closer to the ORF2 region (Fig. 3E, the middle panel). These findings suggest that the repressive effect of the proximal rG4 on ORF2 translation can be attributed to its intrinsic, sequence-specific property, rather than its position relative to the ORF2 region in the transcript. Collectively, our results identified the proximal rG4 as a negative regulator of *MALAT1*-derived sORF2 expression.

### MARS is expressed endogenously in human derived cell line

To determine whether the micropeptide encoded by MALAT1-ORF2 is expressed endogenously in cells, we custom-generated a rabbit polyclonal antibody against the predicted micropeptide. For antigen production, the MALAT1-ORF2 sequence was cloned in-frame with glutathione S-transferase (GST) gene in pGex6P-1 plasmid to express the GST-ORF2 fusion construct in *E. coli* BL21 (DE3) cells (Fig. S3A). The recombinant fusion protein was then affinity-purified (Fig. S3A), and subsequently used as an immunogen to raise anti-sera in two different rabbits. Immunoblotting analyses of uninduced and induced bacterial cell extracts expressing GST-ORF2, and purified GST-ORF2 fusion protein demonstrated that both anti-MARS antisera and the purified anti-MARS antibody exhibited high specificity compared with the pre-immune sera to detect the expected protein (Fig. S3B). These results support the specificity of the generated antibody and its suitability for detecting the endogenous ORF2-derived micropeptide MARS in subsequent cellular analyses.

To examine endogenous MARS expression in U2OS cells, we performed IF studies of U2OS cells, where pre-immune sera and anti-MARS antibody were used to stain those cells. The anti-MARS antibody detected a prominent signal that was predominantly localized to the nucleus, with a lesser cytoplasmic pool, while the pre-immune sera did not produce detectable signal in cells (Fig. 4A). Notably, recent studies have demonstrated that mouse *MALAT1* encodes a micropeptide, M1, which is translated through an internal ribosome entry site (IRES) in neurons (43,44). Alignment of human *MALAT1* with the corresponding region of mouse *MALAT1* revealed modest sequence similarity (Fig. S4A), consistent with previous findings (43). However, the nucleotide sequence corresponding to human MALAT1-ORF2 is not conserved in mouse *MALAT1*. We therefore examined whether MARS is also expressed in mouse cells. IF staining of mouse NIH 3T3 fibroblast cells using the anti-MARS antibody did not show any signal (data not shown). In contrast, the antibody readily detected MARS expression in other human cell types, including retinal pigment epithelial (RPE1) cells and human keratinocytes (HaCaT) (Fig. 4B). These findings suggest that MARS expression is specific to humans.

**Figure 4.**
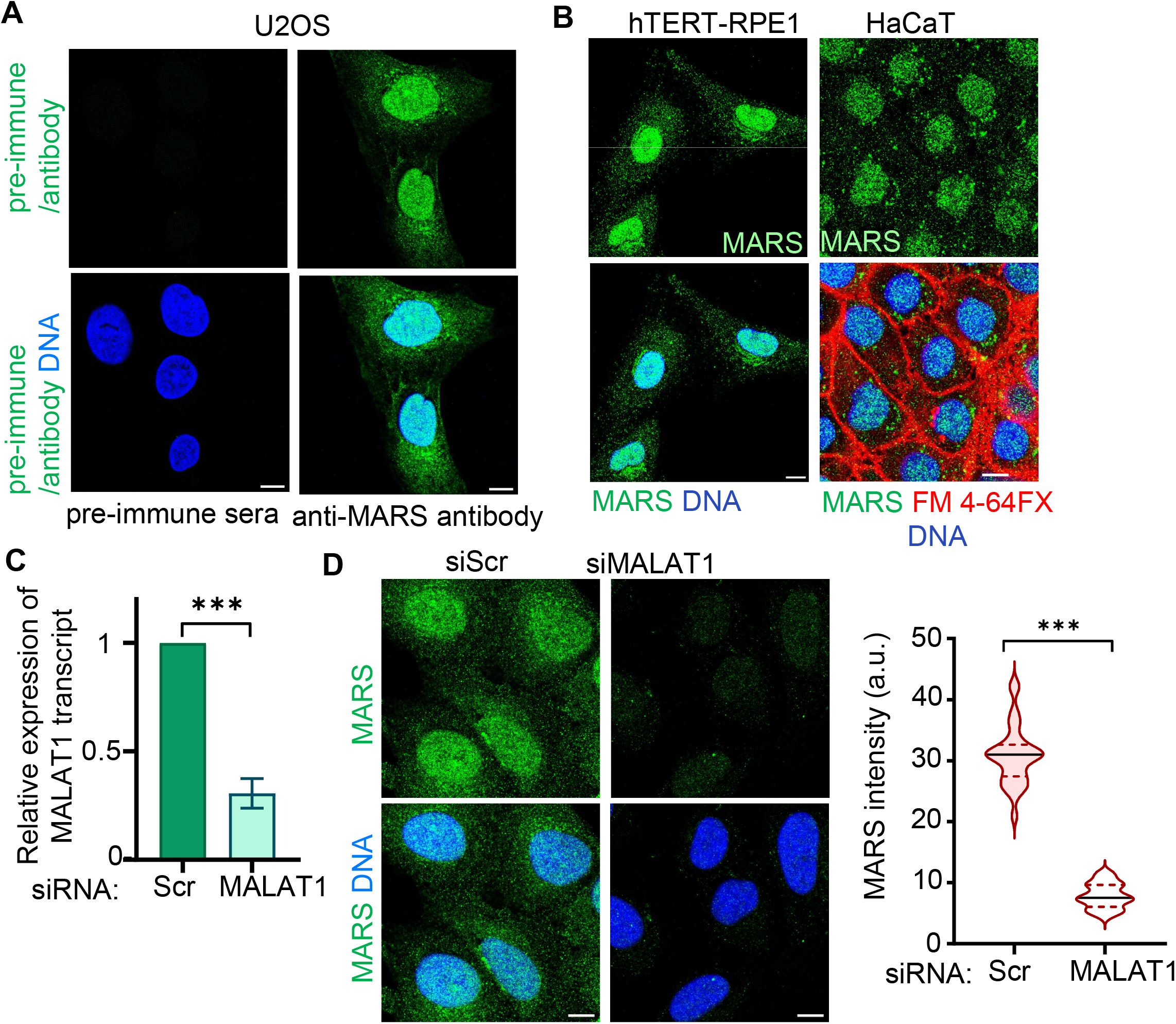
Detection of endogenous expression of MARS in U2OS cell. (A) Immunofluorescence detection of endogenous MARS expression in U2OS cells. Cells were stained with pre-immune serum or anti-MARS primary antibody, followed by the appropriate Alexa fluor tagged secondary antibody. and (B) Detection of MARS in hTERT-RPE1 and HaCaT cells by immunofluorescence using the anti-MARS antibody. MARS staining is shown in green, and the DNA is stained with DAPI (blue). In HaCaT cells, fluorescent probe FM 4-64FX, shown in red, is used to mark the cell boundaries. (C) Relative expression of the *MALAT1* transcript in U2OS cells treated with scrambled (siScr) or *MALAT1*-targeting siRNA (siMALAT1). (D) Representative immunofluorescence images showing MARS expression in siScr and siMALAT1. MARS is shown in green and nuclear DNA in blue. Quantification of background corrected cellular MARS fluorescence intensity is shown on the right. Statistical significance is indicated by asterisks *P < 0.05; **P < 0.01; ***P < 0.001 and ns represents non-significant. The scale bar corresponds to 10 μm.

To establish whether MARS is derived from *MALAT1*, we depleted *MALAT1* transcript in U2OS cells using a *MALAT1*-specific siRNA. *MALAT1* expression was reduced by approximately 75-80% in treated cells (siMALAT1), compared to cells treated with a control scrambled siRNA (siScr; Fig. 4C). Consistent with the proposed origin of the micropeptide, *MALAT1* depletion resulted in a marked reduction in the endogenous MARS signal, as determined by quantitative IF analyses of background-corrected total cellular fluorescence intensities (Fig. 4D). The reduction in endogenous MARS signal following *MALAT1* depletion confirms that expression of MARS depends on the *MALAT1* transcript in U2OS cells, and validates the specificity of the custom-generated anti-MARS antibody.

### Decay from 5’ end of *MALAT1* transcript is required to generate the ORF2 derived peptide, MARS

To determine whether the generation of the MALAT1-derived micropeptide requires 5’ decay of the *MALAT1* transcript, we perturbed the decay pathway by depleting Xrn1 in U2OS cells. Xrn1 is the major cytoplasmic 5′-3′ exoribonuclease in mammalian cells and plays a central role in the degradation of uncapped RNA molecules (4). Given that the *MALAT1* decay intermediate serves as a favourable template for ORF2 translation (Fig. 2), we hypothesized that disrupting XRN1-mediated decay would reduce the abundance of the *MALAT1* decay intermediate and, consequently, decrease MARS expression. Accordingly, we treated U2OS cells with specific siRNA against *XRN1* mRNA, which was used in our previous studies (16,17). As expected, we achieved more than 80% depletion of Xrn1 in siXrn1-treated cells, compared to the control cells (Fig. 5A). Quantitative IF analysis revealed a marked decrease in MARS levels in Xrn1-depleted cells compared with control cells (Fig. 5B). To assess the effect of Xrn1 depletion on *MALAT1* processing, the relative abundance of the decay intermediate and full-length MALAT1 transcript were determined by absolute quantification of transcripts using strategically designed primer pairs that distinguish between the two transcript forms (Fig. 5C). Quantitative PCR revealed a significant reduction in the ratio of the *MALAT1* decay intermediate to the full-length transcript following Xrn1 depletion, indicating reduced accumulation of the decay intermediate under these conditions (Fig. 5D). The reduction in MARS expression, together with the observed decrease in the relative abundance of the decay intermediate upon Xrn1 depletion, suggests that formation of the *MALAT1* decay intermediate is necessary for efficient production of the MALAT1-ORF2-derived micropeptide, MARS. Collectively, these findings support a model in which RNA decay is a prerequisite step in generating a translationally competent *MALAT1* intermediate that serves as a template for production of the micropeptide, MARS.

**Figure 5.**
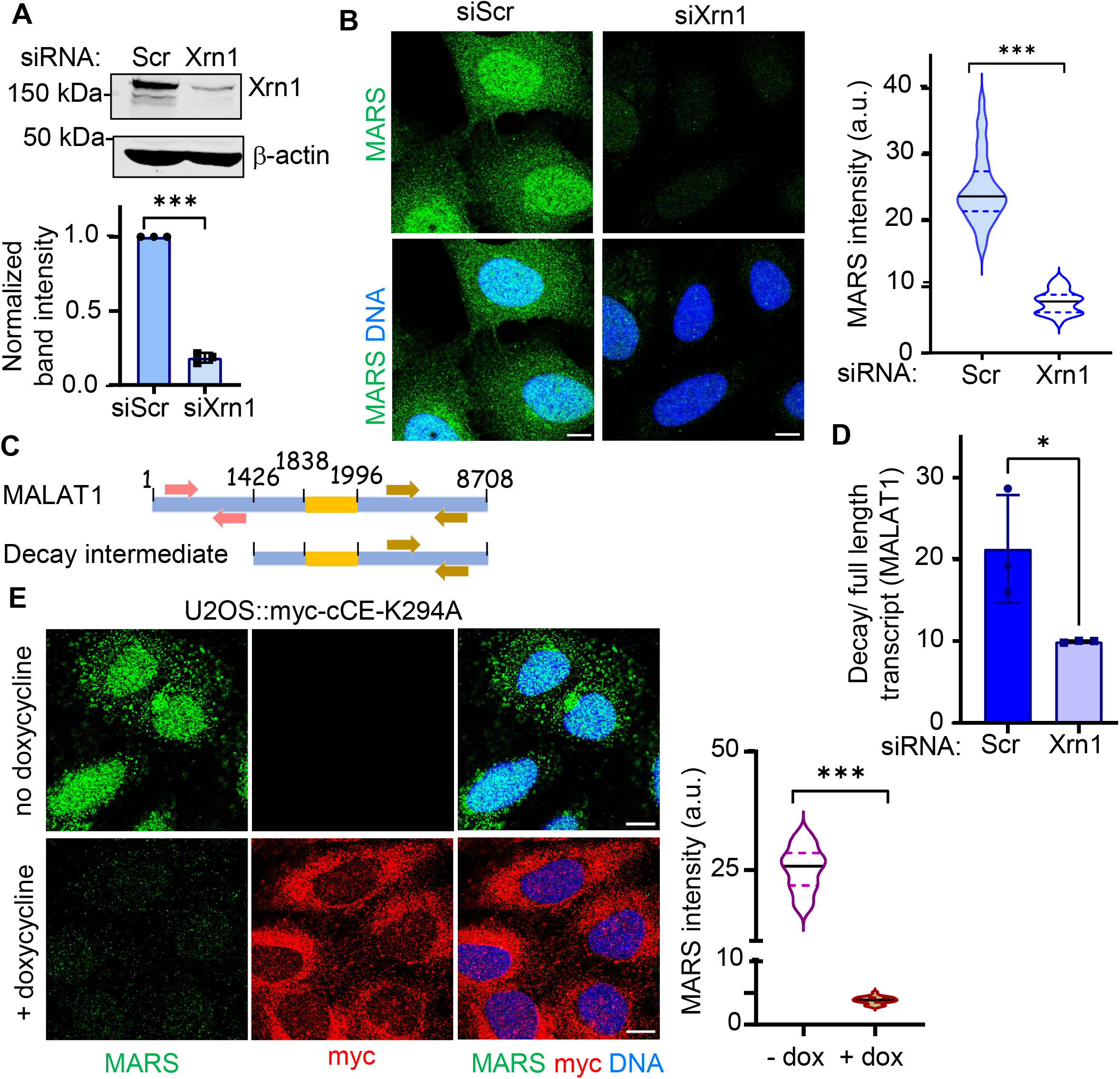
Generation of MARS from MALAT1 depends on cytoplasmic decay and recapping. (A) Immunoblot analysis of Xrn1 knockdown in U2OS cells treated with scrambled siRNA (siScr) or Xrn1-targeting siRNA (siXrn1). The upper panel shows Xrn1 protein levels, and the lower panel shows the corresponding quantification normalized to the loading control β-actin. Molecular weights of the proteins are indicated on the left. (B) Representative immunofluorescence images showing MARS expression following Xrn1 depletion. MARS is shown in green and nuclear DNA in blue, with the corresponding quantification of cellular fluorescence intensity after correcting the background shown on the right. (C) Schematic representation of the full length *MALAT1* transcript and the decay intermediate indicating the positions of the regions used for determining the copy numbers of these two forms of *MALAT1*. (D) Quantification of full-length *MALAT1* and the *MALAT1* decay intermediate by absolute quantification in U2OS cells treated with siScr or siXrn1. XRN1 depletion reduces the ratio of the decay intermediate to full-length MALAT1. (E) Representative immunofluorescence images showing reduced expression of MARS in U2OS cells expressing myc-tagged cCE-K294A in presence of doxycycline. MARS staining is shown in green, myc in red, and nuclear DNA in blue. Quantification of background corrected cellular MARS fluorescence intensity is shown on the right. Statistical significance is indicated by asterisks (*P < 0.05; **P < 0.01; ***P < 0.001) and ns represents non-significant. The scale bar is 10 μm.

### Cytoplasmic recapping is required to generate the ORF2 derived peptide, MARS

We next investigated how MARS is generated from the *MALAT1* decay intermediate. Since the *MALAT1* decay intermediate serves as the template for ORF2 translation, and because expression of ORF2-GFP fusion protein depends on cap-mediated translation (Fig. 2), we hypothesized that cytoplasmic recapping restores the 5’ cap structure of the *MALAT1* decay intermediate, thereby enabling its re-entry into the translational pool.

To test this hypothesis, we utilized our previously employed tetracycline-inducible U2OS cell line expressing myc-tagged K294A, the dominant negative form of capping enzyme, exclusively in the cytoplasm (8,17). As expected, myc-K294A expressed predominantly in the cytoplasm of these cells upon doxycycline (dox) treatment (Fig. 5E, middle panel). Inhibition of cytoplasmic recapping resulted in a marked decrease in the expression of endogenous MARS (Fig. 5E). This reduction suggests that cytoplasmic recapping is required for the production of MARS. Also, this supports our hypothesis that restoration of the 5′ cap structure of the *MALAT1* decay intermediate provides translational competence. Collectively, these findings support a model in which *MALAT1* transcript undergoes a coordinated cycle of 5’ decay and recapping by cytoplasmic activity of CE, generating a recapped decay intermediate that can serve as a template for ORF2 translation.

### MARS exhibits dynamic expression in response to arsenite stress

Previous work from our group demonstrated that cCE targeted transcripts undergo decapping during brief arsenite-induced oxidative stress, followed by recapping during cellular recovery (10). Given that *MALAT1* is regulated by cCE and 5’ decay of *MALAT1*, accompanied by the removal of inhibitory rG4 structures allowing the translation of ORF2 generating MARS, we hypothesized that MARS expression would be regulated in response to oxidative stress.

To test this hypothesis, we treated U2OS cells with 0.5 mM sodium arsenite for 1 h and fixed the cells at 15 min intervals. Cells were stained for MARS and eIF3b, a stress granule (SG) marker to examine the localization of MARS during stress response. Our IF studies revealed colocalization of MARS with SGs (Fig. 6A). Notably, quantitative analyses of total cellular MARS fluorescence showed increased MARS expression level at early time points, reaching a peak approximately 30 min after arsenite treatment, followed by a decline at the later time points (Fig. 6B). Upon recovery from arsenite stress, MARS expression returned to levels comparable to those observed under unstressed conditions (Fig. 6B).

**Figure 6.**
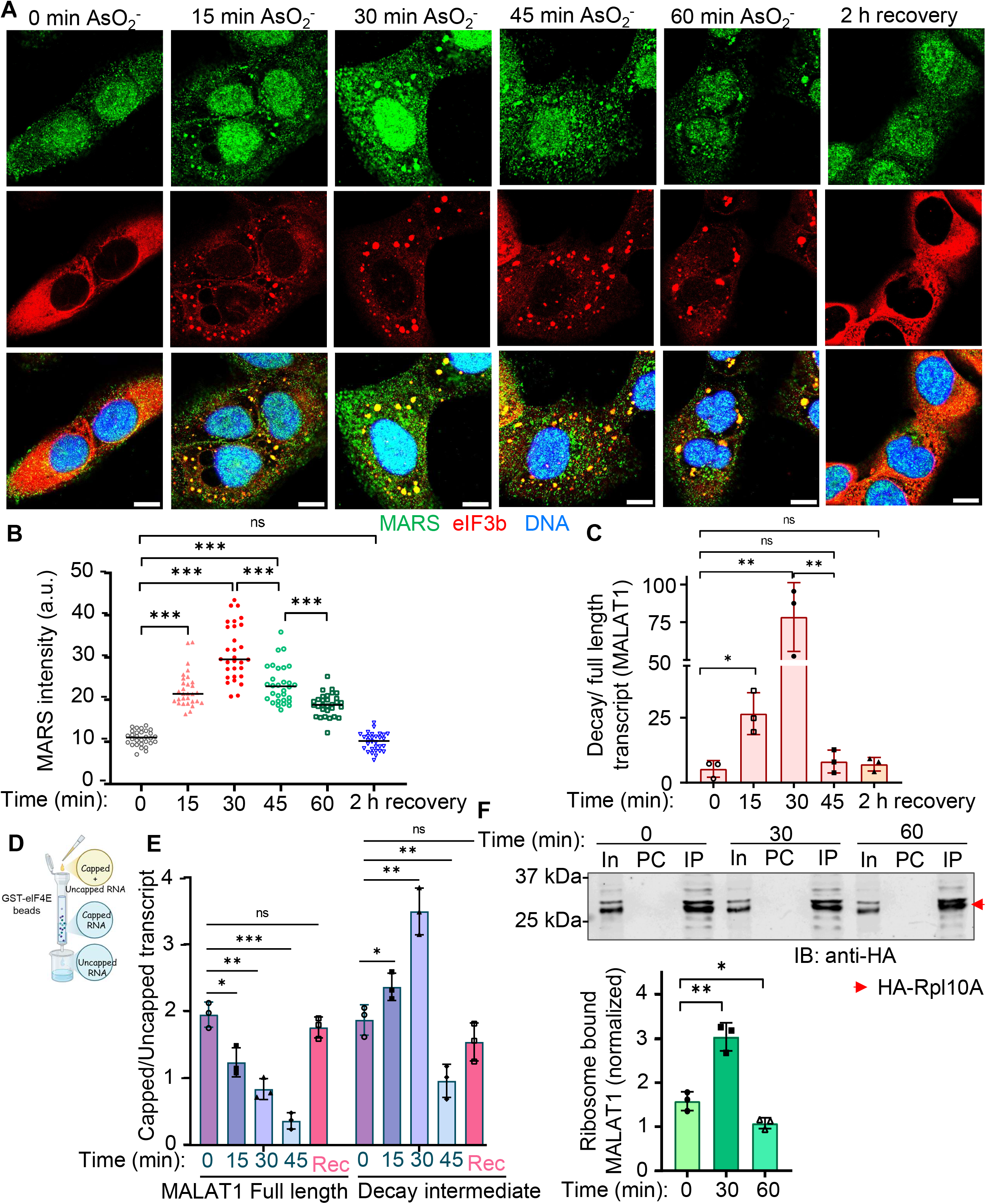
Cellular expression of MARS varies in response to arsenite induced oxidative stress. (A) Representative micrographs showing the distribution of MARS during arsenic stress and subsequent recovery. Cells were treated with 0.05 M of AsO₂⁻ and examined at 0, 15, 30, 45 and 60 min after treatment, followed by analysis after 2 h of recovery. MARS fluorescence increased during the early phase of arsenic exposure, reached its highest level around 30 min, and subsequently decreased, approaching the basal level after recovery. MARS is shown in green, the indicated stress granules marker eIF3b in red, and nuclear DNA in blue. The merged images show colocalization of MARS with eIF3B. The scale bar corresponds to 10 μm. (B) Quantification of MARS fluorescence intensity at the indicated time points following AsO₂⁻ treatment and during recovery (n≥30). (C) Quantification of the *MALAT1* decay-intermediate/full-length transcript ratio at the indicated time points following AsO₂⁻ treatment. The abundance of the decay intermediate increased during the early phase of treatment, with the highest level observed at 30 min, and declined during the later time points and recovery (D) Schematic showing the approach used to separate capped and uncapped transcripts with GST-eIF4E (K119A) immobilized bead as described in earlier study (8). (E) Relative abundance of capped and uncapped full-length *MALAT1* and the *MALAT1* decay intermediate following AsO₂^-^ treatment and recovery. Brief arsenite exposure increased the abundance of the capped *MALAT1* decay intermediate, coinciding with a progressive decrease in capped full-length *MALAT1* abundance up to 30 min following arsenite exposure. (F) Translating ribosome affinity purification (TRAP) was performed at 0, 30 and 60 min following arsenite treatment in cells expressing HA-tagged RPL10A. HA immunoblotting of the input (In), pre-cleared control (PC) and immunoprecipitated (IP) fractions is shown in the upper panel, confirming recovery of HA-tagged RPL10A in the IP fractions at the indicated time points. The lower panel shows the corresponding quantification of ribosome-associated *MALAT1*, normalized to the input *MALAT1*. *MALAT1* association with ribosomes increased at 30 min following arsenite treatment and declined by 60 min. Statistical significance is indicated by asterisks; *P < 0.05, **P < 0.01, ***P < 0.001; ns, not significant.

Although we anticipated that MARS expression would increase during arsenite treatment, MARS abundance decreased at later time points of arsenite-induced stress response. This rapid and dynamic modulation suggests that MARS expression is tightly regulated during the early phases of the cellular stress response, and that its transient nature further indicates a possible role for this *MALAT1*-derived micropeptide in adaptive mechanisms activated shortly after stress exposure. In contrast, when cells were treated with cobalt chloride (CoCl2) to induce hypoxic stress similar to our previous study (17), MARS expression was moderately reduced compared with control cells (Fig. S4B). These findings suggest MARS expression is differentially regulated in response to distinct stress conditions, and may have a potential role in cellular response to arsenite-induced oxidative stress.

To further investigate the mechanism underlying stress-induced expression of MARS and its alterations, we examined the dynamics of full-length and the decay intermediate forms of *MALAT1* during the early phase of arsenite exposure. U2OS cells were treated with 0.5 mM sodium arsenite for varying durations, and the relative abundance of the *MALAT1* decay intermediate and full-length transcript was determined using the quantitative approach described above (Fig. 6C). Such analysis revealed a rapid accumulation of the *MALAT1* decay intermediate during the first 30 minutes of arsenite exposure, indicating enhanced processing of *MALAT1* transcript during the early stress response. This ratio reduced after the 30 min time point of arsenite-induced stress (Fig. 6C). During recovery from arsenite-induced stress, the relative abundance of the *MALAT1* decay intermediate over the full-length transcript returned to levels comparable to those observed in untreated cells (Fig. 6C), which was also observed in the case of MARS expression. These findings suggest that dynamic regulation of *MALAT1* processing is associated with changes in MARS abundance during arsenite-induced stress exposure and recovery, potentially reflecting a role of cap homeostasis in regulating *MALAT1* transcript.

Given our earlier findings that MARS production from the *MALAT1* decay intermediate depends on cap-dependent translation (Fig. 5E), we next investigated if the decay intermediate remained capped during arsenite-induced stress. To assess the cap status of the *MALAT1* full length and its decay intermediate transcripts, capped RNAs were selectively enriched using a GST-tagged fusion protein of the mutated cap-binding protein eIF4E (eIF4E-K119A), which was used to separate the capped RNA effectively from uncapped RNA (12). Briefly, GST-eIF4E(K119A) recombinant protein was expressed in *E. coli* and then immobilized on glutathione-sepharose resin. Total RNA isolated from arsenite-treated cells at the indicated time points, and form cells allowed to recover from arsenite stress, was subjected to affinity purification on the GST-eIF4E(K119A) bound beads (Fig. 6D). RNAs retained by the GST-eIF4E(K119A)-bound beads were classified as the capped transcripts, whereas RNAs recovered in the flow-through were considered as the uncapped fraction (Fig. 6D). Quantitative analysis of the capped and uncapped fractions revealed a progressive reduction in the capped fraction of the full-length *MALAT1* transcript during the first 30 minutes of arsenite treatment. In contrast, the *MALAT1* decay intermediate retained a substantial capped population throughout this period, which reduced after 30 minutes of arsenite exposure (Fig. 6E). These findings suggest that arsenite-induced stress promotes decapping and processing of full-length *MALAT1*, whereas the resulting decay intermediate maintains its capped form likely through cytoplasmic recapping, which, in turn, preserves its translational competence. Collectively, these results support a model in which cytoplasmic recapping maintains cap homeostasis of *MALAT1* and its decay intermediate during the early stages of arsenite-induced stress, which allows the production of the *MALAT1*-derived micropeptide MARS.

To further investigate the translational potential of *MALAT1* during arsenite-induced stress, we performed Translating Ribosome Affinity Purification (TRAP) assays based on the protocol described by Heiman et al (40), with minor modifications. Briefly, U2OS cells expressing HA-tagged Ribosomal Protein L10A (RPL10A) were treated with 0.5 mM sodium arsenite for the indicated durations. Ribosome-associated RNAs were isolated from the immunoprecipitated ribosomes bound to HA-RPL10A (Fig. 6F, upper panel). TRAP analysis revealed dynamic changes in association of *MALAT1* transcript with ribosomes following arsenite exposure (Fig. 6F, lower panel). Specifically, ribosome association of *MALAT1* increased substantially within 30 minutes of arsenite treatment, indicating enhanced recruitment of the *MALAT1* transcript to the translational machinery during the early phase of the stress response. However, this increase was transient, as ribosome-association of *MALAT1* declined sharply by 60 minutes after treatment (Fig. 6F, lower panel). These observations are consistent with the temporal changes of MARS expression and support a model in which early stages of arsenite exposure promote *MALAT1* processing and its association with translating ribosomes. Together with the observed accumulation of the *MALAT1* decay intermediate and its capped population, the increased association of *MALAT1* with ribosomes during the early stages of stress is consistent with enhanced translation of the *MALAT1* ORF2 to produce the micropeptide, MARS. The subsequent decline in ribosome occupancy of *MALAT1* may contribute to the reduction of MARS expression at later time points of arsenite exposure.

### MARS interacts with Nrf2

To gain insight into the biological function of the MARS, we sought to identify its potential interacting partners using the *in silico* protein-protein interaction prediction platform DeepGOWeb (22). Among the highest-confidence candidates, Nrf2, a transcription regulator, emerged as a potential interacting protein of MARS (Fig. 7A-B, S5). Also, Gene Ontology analysis of the predicted MARS interactors, performed using g: Profiler (41) or AmiGO II (23), revealed significant enrichment of GO terms associated with DNA binding and transcriptional regulator activity (Fig. S5). Nrf2 is a master transcriptional regulator of the cellular oxidative stress response and plays a central role in maintaining redox homeostasis (42). Under oxidative stress, Nrf2 activates the transcription of a broad range of cytoprotective and antioxidant genes by binding to AREs present within their regulatory regions (43).

**Figure 7.**
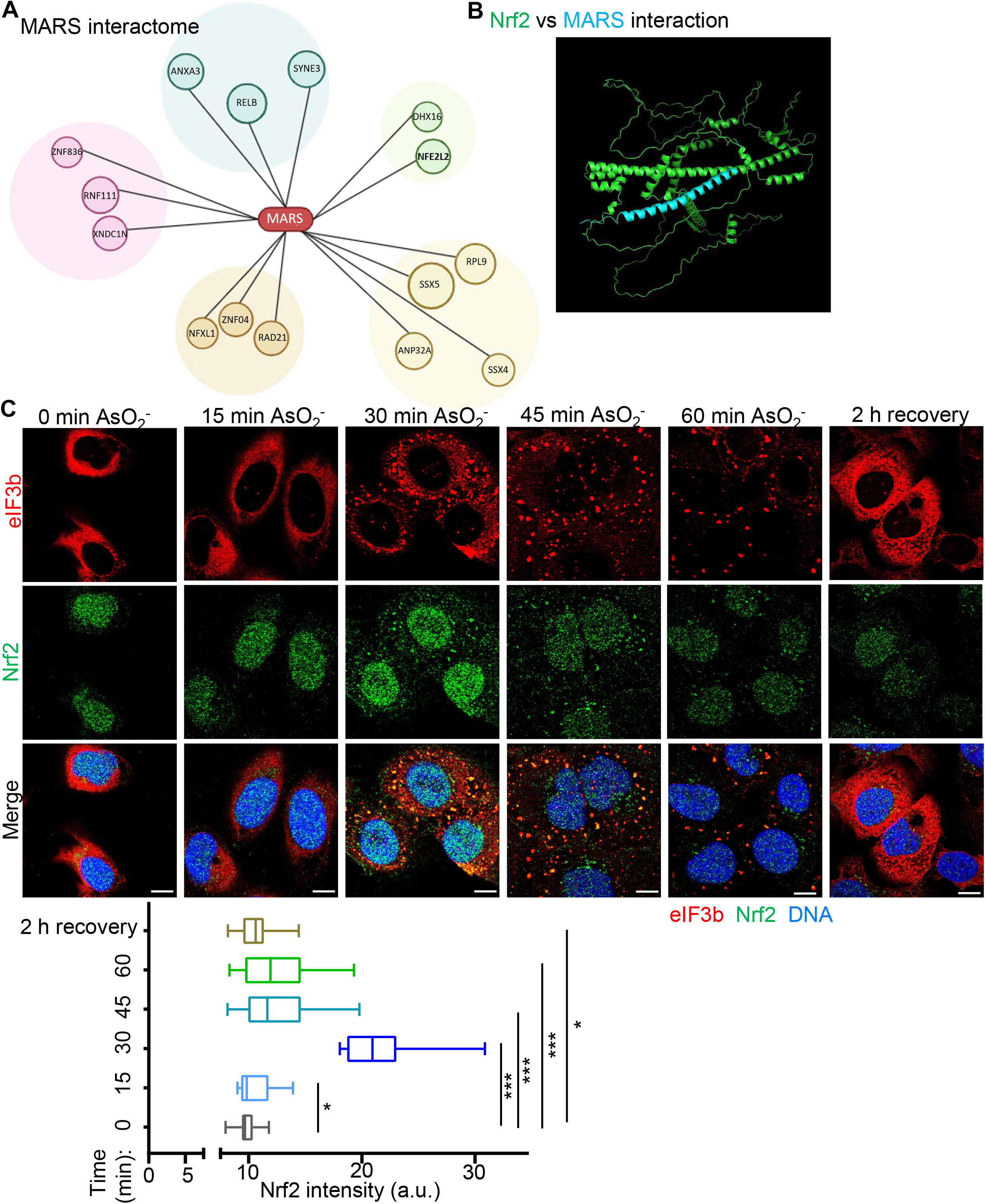
MARS may interact with Nrf2, which exhibits dynamic expression during arsenite induced oxidative stress. (A) Network representation of the proteins identified in the MARS interactome by DeepGOWeb tool. MARS is positioned at the centre of the network, with the identified interacting proteins shown around it. Proteins belonging to related functional groups are represented in the same colour. (B) Predicted interaction between MARS and Nrf2 based on molecular docking analysis by HADDOCK web server. The docked structure of the MARS-Nrf2 complex is shown as a molecular surface representation, with the MARS peptide highlighted in cyan and Nrf2 in green. (C) Representative micrographs showing the distribution of Nrf2 during AsO₂^-^ treatment and recovery. Cells were analysed at 0, 15, 30, 45 and 60 min following AsO₂^-^ treatment and after 2 h of recovery. Nrf2 is shown in green, stress marker eIF3b in red and nuclear DNA in blue. The merged images show the colocalization of Nrf2 with eIF3b marked SGs. Quantification of total Nrf2 fluorescence intensity after background correction at the indicated time points is shown below (n≥50). Nrf2 fluorescence is increased during brief arsenic exposure, reached its highest level around 30 min, and subsequently decreased, approaching the basal level after recovery. Statistical significance is indicated by asterisks; *P < 0.05, **P < 0.01, ***P < 0.001; ns, not significant. The scale bar corresponds to 10 μm.

Protein-protein molecular docking analysis using HADDOCK supported a potentially strong interaction between MARS and Nrf2 (Fig. S6). Given the established role of Nrf2 in coordinating adaptive responses to oxidative stress, its predicted interaction with MARS is particularly intriguing in the context of arsenite-induced stress, during which MARS expression is transiently upregulated. We therefore investigated whether Nrf2 exhibits a temporal expression pattern similar to that of MARS following arsenite exposure. Cells were examined at similar time points during stress or recovery to assess the expression and subcellular localization of Nrf2. Our results showed that Nrf2 partially co-localized with SGs at 30 and 45 min after arsenite exposure, whereas the granular distribution of Nrf2 disappeared at 1 h (Fig. 7C). Quantitative IF analyses revealed that cellular Nrf2 expression level also followed a temporal pattern similar to that of MARS, MARS, reaching its maximum at 30 min after arsenite exposure, and declining at later time points (Fig. 7C). The similar expression patterns of Nrf2 and MARS during oxidative stress and recovery suggests a potential mechanistic link between MARS and oxidative stress signalling pathways.

To experimentally validate the predicted interaction between the MARS and Nrf2, we performed co-immunoprecipitation assays in U2OS cells transiently expressing either EGFP alone or a MARS-EGFP from the ORF2-ATT-GFP fusion construct. Cells were exposed to 0.5 mM sodium arsenite for 30 minutes to induce oxidative stress, after which cell lysates were subjected to immunoprecipitation using an anti-GFP antibody. The resulting immunoprecipitates were subsequently analyzed by immunoblotting for endogenous Nrf2. Co-immunoprecipitation analysis revealed robust enrichment of Nrf2 in complexes isolated from cells expressing the MARS-EGFP fusion protein, whereas Nrf2 was not detected in the GFP-only immunoprecipitates (Fig. 8A). These findings support a specific association between MARS and Nrf2 under arsenite-induced stress conditions. The stress-dependent interaction is consistent with the transient induction of MARS expression following arsenite exposure and suggests a potential role of MARS in oxidative stress signalling. Collectively, these observations raise the possibility that MARS modulates Nrf2-mediated adaptive responses during the early stages of cellular stress, potentially influencing the expression of antioxidant defence genes.

**Figure 8.**
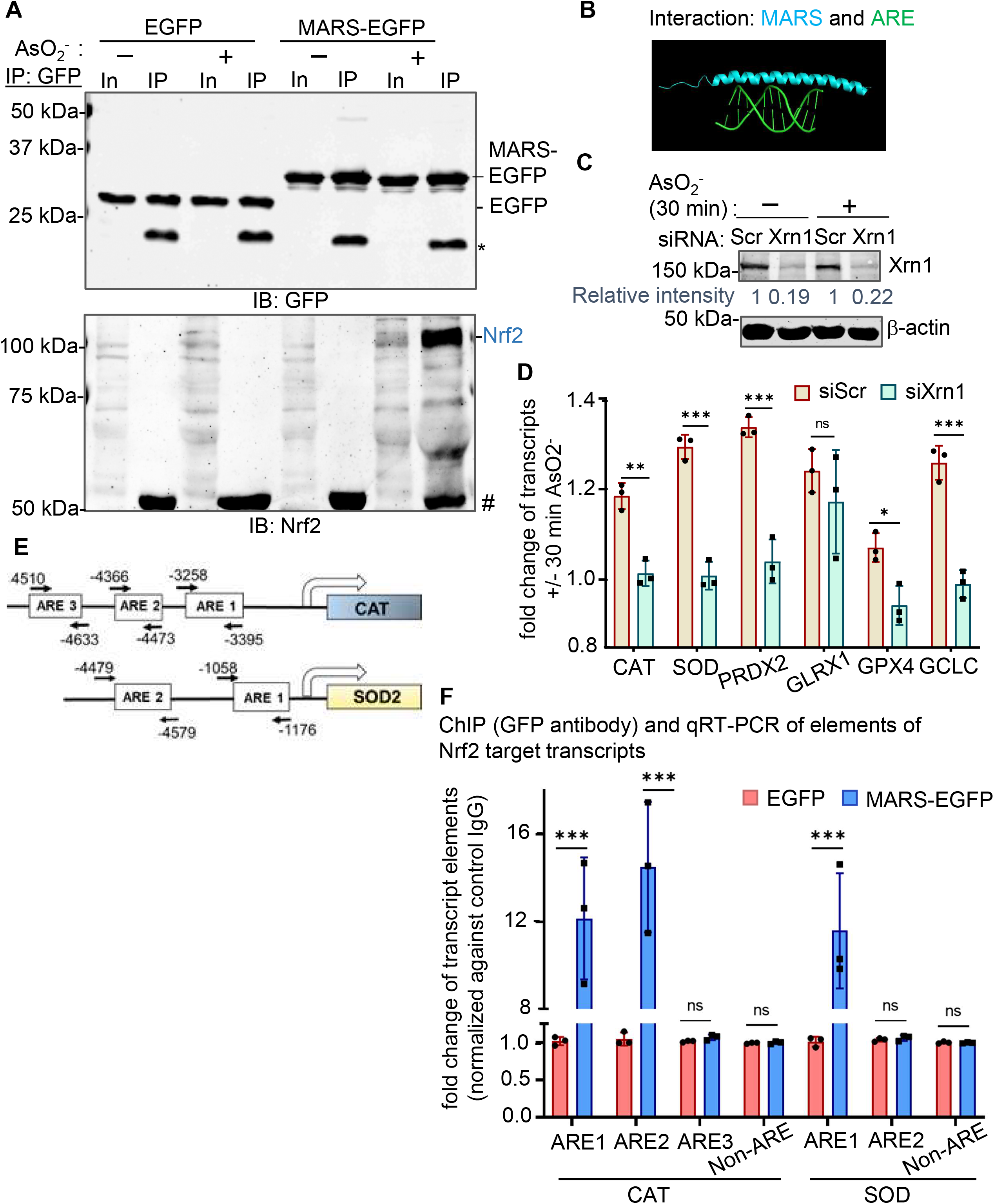
MARS associates with Nrf2 and genes containing ARE. (A) Co-immunoprecipitation analysis to examine the association of MARS with Nrf2. Cells expressing either EGFP or MARS-EGFP were subjected to immunoprecipitation using an anti-GFP antibody after a brief exposure of AsO₂^-^ for 30 min or without AsO₂^-^ treatment, and the immunoprecipitated proteins were analysed by immunoblotting for GFP and Nrf2. Nrf2 was detected in the MARS-GFP immunoprecipitate but not in the GFP control after AsO₂^-^ exposure. (B) Molecular docking model showing the predicted interaction of MARS with the ARE containing DNA. (C) Cells were treated with either control (siScr) or Xrn1-targeting siRNA (siXrn1) and exposed to AsO₂^-^ for 30 min or kept untreated. The blot demonstrates the successful knock-down of Xrn1 in siXrn1 treated samples compared to siScr treated controls. Quantification of Xrn1 knockdown efficiency was performed using ImageJ software, with β-Actin serving as the internal loading control. (D) The panel shows the reduced expressions of ARE containing candidate genes in siXrn1 treated samples with reference to siScr treated samples during 30 mins of arsenite treatment compared to untreated cells as measured by quantitative real-time PCR. (E) The promoter organization of the CAT and SOD2 genes is shown below, indicating the positions of the predicted AREs relative to the transcription start site. (F) ChIP was performed using an anti-GFP antibody, followed by qRT-PCR to assess the enrichment of selected antioxidant response element (ARE)-containing regions of the *CAT* and *SOD2* promoters in cells expressing MARS-EGFP. Cells expressing GFP alone served as the control. Enrichment was normalized to the IgG control, and regions lacking ARE sequences (Non-ARE) were included as negative controls. Data are presented as fold change relative to the GFP control. Statistical significance is indicated as follows: ns, not significant; *P* < 0.05; P < 0.01; *P* < 0.001.

### MARS may act as a transcription regulator of ARE containing dsDNAs

To investigate whether MARS contributes to Nrf2-mediated transcriptional responses during oxidative stress, we examined its potential interaction with ARE containing DNA. Molecular docking analysis using HADDOCK predicted a favourable interaction between MARS and ARE containing double stranded (ds) DNA (Figs. 8B and S7). In contrast, however, the predicted interaction between MARS and or RNA (data not shown) or MARS and hypoxia response element (HRE)-containing dsDNA was less favourable, as indicated by the HADDOCK score (Fig. S7). These findings suggest that MARS may preferentially interact with ARE-containing DNA. However, to validate this speculation and to test how MARS influences the expression of ARE containing DNAs, expressions of canonical ARE containing genes under conditions that do not support MARS expression were determined.

As described above, MARS expression requires 5’ end decay of *MALAT1*, followed by translation of the recapped decay intermediate. MARS production was disrupted by depleting Xrn1, the major mammalian 5′-3′ exoribonuclease, whose activity is required for accumulation of the *MALAT1* decay intermediate and subsequent translation of MALAT1-ORF2 (Fig. 5). The same Xrn1 depletion strategy was employed here (Fig. 8C). Following Xrn1 depletion, U2OS cells were exposed to 0.5 mM sodium arsenite for 30 minutes, and the expression of representative Nrf2-responsive, ARE-containing genes such as Catalase (CAT), Superoxide Dismutase 2(SOD2), Peroxiredoxin 2 (PRDX2), Glutaredoxin 1 (GLRX1), Glutathione peroxidase 4 (GPX4), and Glutamate-cysteine ligase catalytic subunit (GCLC) was assessed. Compared with control cells, Xrn1 depleted cells exhibited significantly reduced induction of CAT, SOD2, PRDX2, GLRX1, GPX4, and GCLC (Fig. 8D). The reduced expression of these antioxidant genes suggests that Xrn1 depletion compromises the cellular response to arsenite-induced oxidative stress, likely by reducing MARS biogenesis. These findings are consistent with the possibility that MARS contributes to the regulation of Nrf2-responsive genes during arsenite-induced stress. However, because Xrn1 depletion may also affect other RNA-processing pathways, the observed changes cannot be attributed exclusively to impaired MARS production.

To investigate whether MARS associates with the ARE regions of Nrf2-responsive genes, we examined the promoter regions of CAT and SOD2 genes, and designed primers targeting three putative ARE-containing regions in the CAT promoter and two in the SOD2 promoter (Fig. 8E). Primers targeting regions outside the predicted ARE sequences (non-ARE) in the promoter region of corresponding genes were also included as negative controls. We performed chromatin immunoprecipitation (ChIP) assays in U2OS cells expressing either EGFP alone or MARS-EGFP fusion protein. Chromatin-associated complexes were immunoprecipitated using an anti-GFP antibody, followed by quantitative PCR analysis using primers described above (Fig. 8E). ChIP-qPCR analysis revealed significant enrichment of the ARE-containing regions (ARE1 and ARE2 of the CAT gene and ARE1 of the SOD2 gene promoter) in chromatin fractions immunoprecipitated from cells expressing the MARS-EGFP fusion protein (Fig. 8F). In contrast, no appreciable enrichment was detected in cells expressing GFP alone, indicating that the observed promoter association is specific to MARS. Notably, the binding of MARS to the promoter sequences of either CAT or SOD2 was not uniform across all predicted ARE-containing regions, since neither CAT ARE3 not SOD2 ARE2 showed appreciable enrichment (Fig. 8F). This differential enrichment suggests that the association of MARS-containing complexes with promoter regions of AGE containing genes may vary depending on the cellular context. Nonetheless, the enrichment of selected ARE-containing promoter regions suggests that MARS may associate with regulatory regions of Nrf2-responsive genes. This association provides a potential mechanistic link between MARS expression during the early time of arsenite-induced oxidative stress and the regulation of antioxidant defence programs. However, further investigation is required to determine whether MARS directly interacts with these ARE-containing promoter regions or is recruited to these regions through interactions with other chromatin-associated proteins, which is beyond the scope of this study. Additional studies will also be necessary to establish the extent to which MARS contributes directly to the transcriptional regulation of these genes.

## Discussion

Previous studies have established a role for cytoplasmic capping in regulating transcript stability through modulation of the 5′ cap status. Most of these studies examined the cap status of transcripts under stress conditions, such as arsenite-induced stress (10) or CoCl2-induced hypoxia (17), in cells with active or inactive cytoplasmic capping activity. These studies proposed that the cytoplasmic capping enzyme (cCE) restores 5’ cap on its target transcripts following decapping (10,17). It was further hypothesized that recapped transcripts may re-enter the translational pool and may generate new proteins (8). This model was supported by the observation that uncapped transcripts and their wild-type counterparts exhibited similar poly(A)-tail lengths, suggesting that recapping may be necessary to restore cap-dependent translation (44).

Advances in ribosome profiling and other high-resolution techniques recently revealed that numerous sORFs encode previously unrecognized micropeptides with potential regulatory functions (45–49). These findings prompted us to investigate whether cCE-regulated lncRNAs could similarly serve as templates for micropeptide production through ORFs embedded within their sequences. Our previous study identified stable, uncapped decay intermediates derived from the cCE-regulated transcripts *MALAT1, GAS5-838, RP1-283E3.8*, and *NFYC-AS1* (15). These decay intermediates accumulated in cells inducibly expressing the cytoplasmic restricted, catalytically inactive mutant of CE (K294A) (15). We had also sequenced these decay intermediates to define their uncapped ends (15). In the present study, we reanalysed these sequences and identified multiple ORFs, along with evidence of their ribosome association. When transiently expressed in U2OS cells using a chimeric GFP construct, all identified ORFs produced GFP-fusion proteins, as confirmed by fluorescence microscopy and immunoblotting (Fig. 1). These findings support the coding potential of the identified ORFs and demonstrate their ability to generate peptides in the reporter system. These micropeptides exhibited distinct subcellular localizations, both among peptides derived from the same parental lncRNA, such as ORF1 and ORF2 of *MALAT1*, and among peptides derived from different lncRNAs, such as ORF1 of *GAS5-838* and ORF2 of *RP1*. These expression and localization patterns suggest that the putative micropeptides may have distinct biological properties rather than representing nonspecific by-products of lncRNA translation.

Among these candidates, we selected *MALAT1* ORF2 for further investigation because it overlapped with a ribosome-protected fragment, and was recovered within a *MALAT1* decay intermediate identified by 5′-RACE cloning in our previous study (15). Notably, reporter GFP expression was detected when either the ORF2 or the decay intermediate containing ORF2 was cloned into the chimeric reporter construct. In contrast, GFP expression was not observed when the entire 5′ region of *MALAT1* containing ORF2 was used (Fig. 2). These findings suggest that the 5′ region of *MALAT1* imposes a constraint on ORF2 translation. Controlled degradation of this 5′ region may relieve the inhibitory effect, enabling the resulting decay intermediate to undergo cap-dependent translation and produce MALAT1-derived MARS micropeptide. Thus, our findings support a model in which cytoplasmic processing of *MALAT1* generates a translationally-competent decay intermediate that serves as a template for MARS production.

To investigate the potential basis of translational repression of MALAT1*-*ORF2, which might be embedded in the 5’ region of *MALAT1*, we examined the sequence of this region for rG4 motifs. Multiple rG4s have previously been identified within *MALAT1* (3765–8158), where they contribute to binding with the NONO protein (49). Our analysis identified two new rG4s motifs upstream of ORF2, within 1-1400 nucleotides of *MALAT1* transcript. We further demonstrated that the rG4 motif proximal to ORF2 inhibits the translation of ORF2. Notably, this inhibitory effect depends on the rG4 sequence itself rather than its proximity to ORF2. Relocating the proximal rG4 to a distal position similarly suppressed ORF2 translation, whereas mutating the sequence of this motif to disrupt rG4 formation, irrespective of its original position or after relocation to a distal position, partially restored ORF2 translation (Fig. 3). These findings suggest that the rG4 sequence acts as a cis-regulatory element controlling MARS translation from ORF2. Similar inhibitory effects of rG4 structures on translation have been reported in other transcripts, which were mRNAs (39,48). Interestingly, the 5′ UTR of mouse *MALAT1* exhibits a distinct regulatory architecture, containing stem-loop structures that embed an IRES, which mediate cap-independent translation (50). In contrast, the 5’ region of human *MALAT1* contains inhibitory rG4 motifs that regulate MARS translation. These differences highlight species-specific organization of the *MALAT1* 5′ regions and raise the possibility that different RNA structural elements have evolved to regulate the translation of different *MALAT1*-encoded micropeptides in a species and context-dependent manner.

MALAT1-derived micropeptide MARS was detected in human-derived cell lines, including U2OS, hTERT-RPE1, and HaCaT, but was not detected in NIH 3T3 cells (Fig. 4). This observation is consistent with the limited conservation of the human and mouse *MALAT1* sequences, and further suggests that MARS biogenesis depends on species-specific sequence features. The detection of endogenous MARS in human cells supports the possibility that endogenous *MALAT1* serves as its precursor, potentially following 5′-end decay that removes the inhibitory rG4 motifs. This model is further supported by our previous identification of a stable *MALAT1* decay intermediate containing ORF2 (15), which could serve as a template for MARS translation.

Consistent with this model, depletion of either *MALAT1* or *XRN1* reduced MARS expression, indicating that both the parental RNA and the 5′-to-3′ exonuclease XRN1 are necessary for MARS biogenesis. Our subsequent experiment that examined the abundance of the *MALAT1* decay intermediate relative to that of the full-length transcript demonstrated reduction in the decay intermediate population following Xrn1 depletion, thereby providing potential mechanistic explanation for the decreased MARS expression in these cells. Furthermore, the decay intermediate appears to undergo cytoplasmic recapping, as inhibition of cytoplasmic capping markedly reduced MARS expression (Fig. 5). Together, these findings provide, to our knowledge, the first evidence that cCE-mediated recapping of an uncapped *MALAT1* decay intermediate may facilitate subsequent translation of an ORF embedded in it. Thus, this mechanism supports the concept of “cap homeostasis” of specific transcripts.

In our previous study, we demonstrated that cCE targeted mRNA transcripts lose their 5’ cap during arsenite induced oxidative stress, and regain their cap during recovery (10). We therefore reasoned that if *MALAT1*, being a cCE target, undergoes similar stress-dependent processing, the resulting decay intermediate could provide additional template for MARS translation. Consistent with this possibility, MARS expression exhibited a dynamic pattern during arsenite exposure, with high expression at early time points and lower expression at later time points, although expression at 1 h remained higher than that in untreated cells. Consistent with these observations, the accumulation of the *MALAT1* decay intermediate relative to full-length MALAT1, its recapped state, and ribosome association are significantly enhanced at early time points following arsenite exposure, and declined at later time points (Fig. 6). These temporal changes support a model in which arsenite-induced stress promotes the early accumulation and translational engagement of the MALAT1 decay intermediate, thereby contributing to increased MARS production. Notably, a similar dynamic pattern of MARS expression was not observed under CoCl₂-induced hypoxia, suggesting that MARS expression may be associated more closely with the arsenite-induced oxidative stress response than with the hypoxic conditions examined.

This observation prompted us to investigate the potential protein interactors of MARS using *in silico* prediction tools. Among the candidate interactors, Nrf2 was particularly interesting because of its established role in regulating the antioxidant responses through various ARE-containing genes (42,43,51). Nrf2 regulates several oxidant-responsive cellular programs, including autophagy, inflammasome assembly, endoplasmic reticulum (ER) stress and the unfolded protein response (UPR) (51). The recovery of Nrf2 in MARS-containing immunoprecipitated complex following arsenite exposure (Fig. 7) raises the possibility that MARS indeed associates with Nrf2 and participates in oxidative stress response.

Although Nrf2 biology has been extensively investigated, the precise composition of the Nrf2-associated transcriptional complexes and the regulatory interactions governing their assembly and function remain incompletely understood (52). Under basal conditions, Nrf2 is maintained in low level at cytoplasm since it associates with Kelch-like ECH-associated protein 1 (KEAP1) that promotes its ubiquitination and proteasomal degradation (53,54). Upon oxidative stress, Nrf2 dissociates from KEAP1 and stabilized, translocates to the nucleus, and form heterodimers with small Maf (sMAF) proteins to regulate the transcription of ARE-containing genes (52,54,55). The observed association between MARS and Nrf2 therefore raises the possibility that MARS contributes to the regulation of Nrf2-associated transcriptional responses during oxidative stress.

Our *in silico* modelling and experimental outcomes further suggest that MARS associates with several ARE-containing regions of antioxidant genes, including SOD2 and CAT, and regulates their expression, and that disruption of MARS biogenesis through XRN1 depletion is accompanied by reduced expression of those antioxidant genes (Fig. 8). Together, these observations support a potential role for MARS, in association with Nrf2, in modulating antioxidant defence programmes during arsenite-induced stress. However, the precise mechanism underlying this regulation remains to be established. In particular, further studies are needed to determine whether MARS directly interacts with sMAF proteins or other components of the Nrf2 transcriptional complex and whether such interactions contribute to the regulation of Nrf2-dependent transcription.

Future studies should also investigate the functions of MARS in other cellular contexts in which Nrf2 plays an important role, including inflammation, autophagy, and related stress-response pathways. Determining whether MARS can modulate oxidative stress independently of Nrf2 would provide additional insight into its biological function. Notably, the micropeptide encoded by mouse *MALAT1* has also been implicated in the modulation of synaptic function (56), while another study reported that the human *MALAT1*-derived micropeptide promotes Influenza A virus replication by attenuating the innate immune response in A549 cells (57). These findings further highlight the potential context-dependent functions of MALAT1-derived micropeptide and underscore the importance of investigating MARS in diverse cellular and physiological contexts.

In conclusion, our study identifies a mechanism linking cytoplasmic processing and recapping of a lncRNA, *MALAT1,* to the biogenesis of micropeptide MARS. Our findings reveal a new layer of post-transcriptional regulation, in which cytoplasmic recapping contributes to proteome diversification. The biogenesis of MARS from a stable decay intermediate of *MALAT1* highlights a previously unrecognized mode of gene regulation where the decay intermediate also acts as a precursor for micropeptide production through cytoplasmic processing.

Furthermore, our findings suggest that MARS production is influenced by RNA structural elements and cellular stress and may be associated with the regulation of Nrf2-responsive antioxidant genes during arsenite-induced oxidative stress. Together, cytoplasmic RNA processing and recapping can contribute to micropeptide production from a lncRNA, and that this process may be regulated by RNA structure and cellular stress. Future studies should investigate the biogenesis, regulation and biological function of additional micropeptides derived from other lncRNAs regulated by cytoplasmic recapping.

## Data Availability

All data are contained within the manuscript and will be available upon reasonable request directed to the corresponding author, Chandrama Mukherjee.

## Supplementary Data

This article contains supporting information.

## Acknowledgments

The authors thank Dr. Somsubhra Nath (Presidency University, Kolkata, India), Dr. Benubrata Das (IACS, Kolkata, India) and Prof. Partha Saha (SINP, Kolkata, India) for providing the constructs expressing pEGFP-N1, mcherry-H2B and HCV-IRES. The authors also thank Dr. Debanjan Mukhopadhyay (Presidency University, Kolkata, India), for providing the HaCaT cell line, the Institute of Health Sciences, Presidency University for the departmental instrument facility, and Presidency University for necessary infrastructural support.

## Author contributions

Conceptualization: A.M. and C.M.; Methodology: A.M., R.S., P.P.M and C.M.; Software: A.M. and R.S. Formal analysis: A.M., R.S., P.P.M., S.M. and C.M Investigation: A.M., R.S., P.P.M. and C.M; Resources: P.P.M. and C.M; Data curation: A.M., R.S., P.P.M. and C.M.; Writing – original draft: A.M. and C.M..; Writing – review & editing: A.M., R.S., P.P.M., S.M. and C.M.; Visualization: A.M., R.S., P.P.M., S.M. and C.M.; Supervision: C.M. Project administration: C.M. Funding acquisition: C.M.

