## Supplementary Figures and Table-1 for "Cytoplasmic recapping commences the translation of a micropeptide, MARS from MALAT1 after the removal of an inhibitory RNA G-quadruplex"

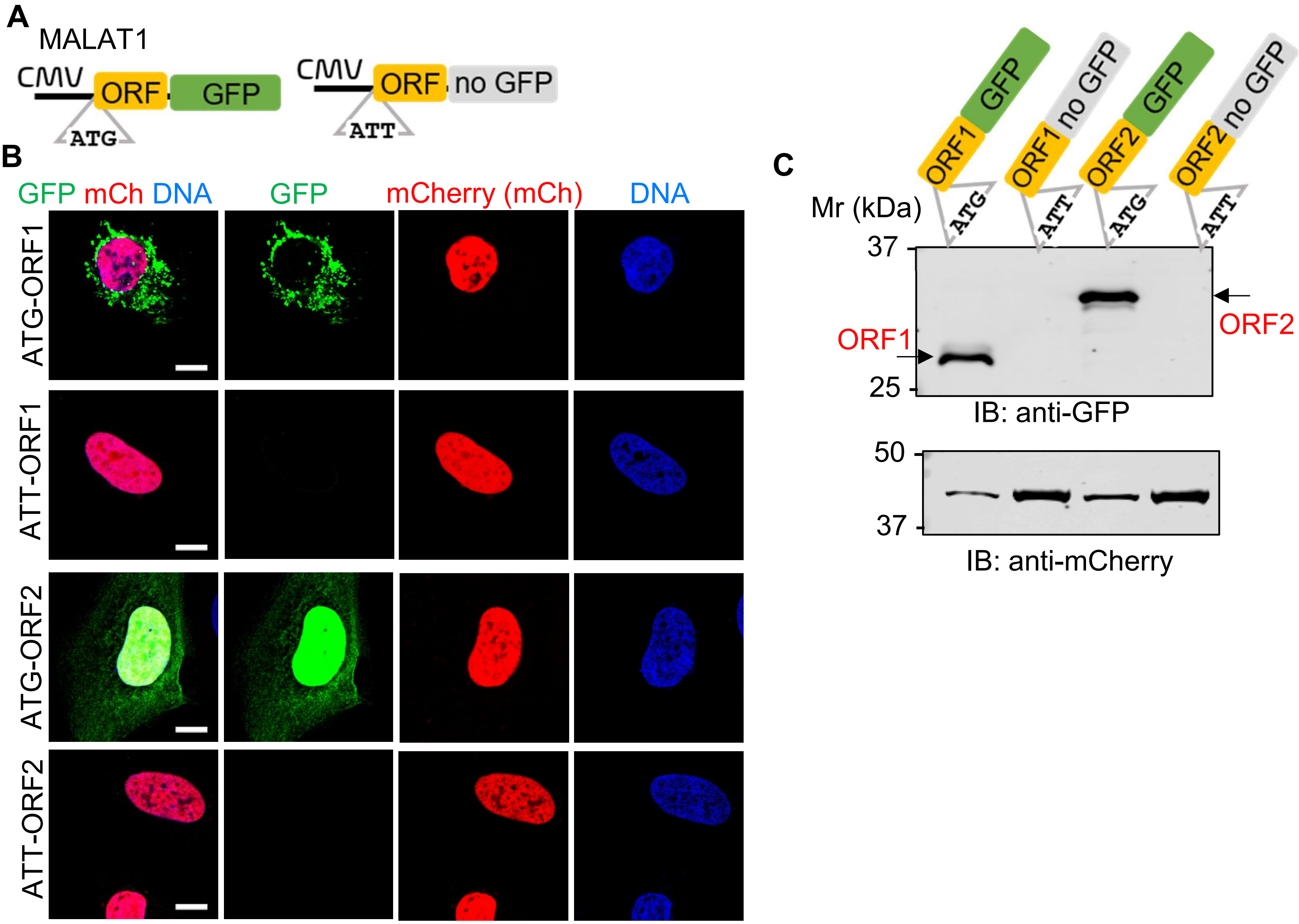


**Supplementary Figure 1. GFP Chimeric protein expression depends on the start codon of ORFs of MALAT1.**

(A) Schematic representation of the reporter constructs used to examine whether the predicted ORFs of *MALAT1* can initiate translation from its start codon. The constructs contain the indicated ORF sequences positioned upstream of mutated pEGFPN1 vector, with either ATG or ATT at the corresponding translation initiation site. (B) Representative fluorescence microscopy images of cells expressing the indicated ATG or ATT containing ORF1 and ORF2-EGFP reporter constructs. Constructs carrying ATG-ORF1 or ATG-ORF2 show GFP fluorescence, whereas replacement of the initiation codon of the candidate ORFs with ATT abolished the GFP signal. mCherry-H2B is used as a control for transfection. Scale bar is 10 µm. (C) Immmunoblot analysis of indicated constructs with anti-GFP antibody to detect the expression of ORF-GFP chimeras. Western hybridization with anti-mcherry antibody shows similar transfection of the constructs in cells. Molecular weight of the proteins is indicated in the left.

ClustalW alignment of the sequences of human MALAT1 transcript (partial), MALAT1 decay intermediate (identified in Mukherjee et al., 2023) and the MALAT1 ORF2 (identified in this study)

```
                   10        20        30        40        50        60        70        80
         ....|....|....|....|....|....|....|....|....|....|....|....|....|....|....|....|
MALAT1   GTAAAGGACTGGGGCCCCGCAACTGGCCTCTCCTGCCCTCTTAAGCGCAGCGCCATTTTAGCAACGCAGAAGCCCGGCGC
5'-RACE  --------------------------------------------------------------------------------
ORF2     --------------------------------------------------------------------------------
.............
                 1370      1380      1390      1400      1410      1420      1430      1440
         ....|....|....|....|....|....|....|....|....|....|....|....|....|....|....|....|
MALAT1   TCTCTTTGAAAGATAGAGATTAATACAACTACTTAAAAAATATAGTCAATAGGTTACTAAGATATTGCTTAGCGTTAAGT
5'-RACE  -----------------------------------------------------------------TGCTTAGCGTTAAGT
ORF2     --------------------------------------------------------------------------------

                 1450      1460      1470      1480      1490      1500      1510      1520
         ....|....|....|....|....|....|....|....|....|....|....|....|....|....|....|....|
MALAT1   TTTTAACGTAATTTTAATAGCTTAAGATTTTAAGAGAAAATATGAAGACTTAGAAGAGTAGCATGAGGAAGGAAAAGATA
5'-RACE  TTTTAACGTAATTTTAATAGCTTAAGATTTTAAGAGAAAATATGAAGACTTAGAAGAGTAGCATGAGGAAGGAAAAGATA
ORF2     --------------------------------------------------------------------------------

                 1530      1540      1550      1560      1570      1580      1590      1600
         ....|....|....|....|....|....|....|....|....|....|....|....|....|....|....|....|
MALAT1   AAAGGTTTCTAAAACATGACGGAGGTTGAGATGAAGCTTCTTCATGGAGTAAAAAATGTATTTAAAAGAAAATTGAGAGA
5'-RACE  AAAGGTTTCTAAAACATGACGGAGGTTGAGATGAAGCTTCTTCATGGAGTAAAAAATGTATTTAAAAGAAAATTGAGAGA
ORF2     --------------------------------------------------------------------------------

                 1610      1620      1630      1640      1650      1660      1670      1680
         ....|....|....|....|....|....|....|....|....|....|....|....|....|....|....|....|
MALAT1   AAGGACTACAGAGCCCCGAATTAATACCAATAGAAGGGCAATGCTTTTAGATTAAAATGAAGGTGACTTAAACAGCTTAA
5'-RACE  AAGGACTACAGAGCCCCGAATTAATACCAATAGAAGGGCAATGCTTTTAGATTAAAATGAAGGTGACTTAAACAGCTTAA
ORF2     --------------------------------------------------------------------------------

                 1690      1700      1710      1720      1730      1740      1750      1760
         ....|....|....|....|....|....|....|....|....|....|....|....|....|....|....|....|
MALAT1   AGTTTAGTTTAAAAGTTGTAGGTGATTAAAATAATTTGAAGGCGATCTTTTAAAAAGAGATTAAACCGAAGGTGATTAAA
5'-RACE  AGTTTAGTTTAAAAGTTGTAGGTGATTAAAATAATTTGAAGGCGATCTTTTAAAAAGAGATTAAACCGAAGGTGATTAAA
ORF2     --------------------------------------------------------------------------------

                 1770      1780      1790      1800      1810      1820      1830      1840
         ....|....|....|....|....|....|....|....|....|....|....|....|....|....|....|....|
MALAT1   AGACCTTGAAATCCATGACGCAGGGAGAATTGCGTCATTTAAAGCCTAGTTAACGCATTTACTAAACGCAGACGAAAATG
5'-RACE  AGACCTTGAAATCCATGACGCAGGGAGAATTGCGTCATTTAAAGCCTAGTTAACGCATTTACTAAACGCAGACGAAAATG
ORF2     -----------------------------------------------------------------------------ATG

                 1850      1860      1870      1880      1890      1900      1910      1920
         ....|....|....|....|....|....|....|....|....|....|....|....|....|....|....|....|
MALAT1   GAAAGATTAATTGGGAGTGGTAGGATGAAACAATTTGGAGAAGATAGAAGTTTGAAGTGGAAAACTGGAAGACAGAAGTA
5'-RACE  GAAAGATTAATTGGGAGTGGTAGGATGAAACAATTTGGAGAAGATAGAAGTTTGAAGTGGAAAACTGGAAGACAGAAGTA
ORF2     GAAAGATTAATTGGGAGTGGTAGGATGAAACAATTTGGAGAAGATAGAAGTTTGAAGTGGAAAACTGGAAGACAGAAGTA

                 1930      1940      1950      1960      1970      1980      1990      2000
         ....|....|....|....|....|....|....|....|....|....|....|....|....|....|....|....|
MALAT1   CGGGAAGGCGAAGAAAAGAATAGAGAAGATAGGGAAATTAGAAGATAAAAACATACTTTTAGAAGAAAAAAGATAAATTT
5'-RACE  CGGGAAGGCGAAGAA-----------------------------------------------------------------
ORF2     CGGGAAGGCGAAGAAAAGAATAGAGAAGATAGGGAAATTAGAAGATAAAAACATACTTTTAGAAGAAAAAAGATAA----
.............
                 2170      2180      2190      2200
         ....|....|....|....|....|....|....|....|
MALAT1   AAGTATTGGAGAAGTATAGAAGATAGAAAAATATAAAGCC
5'-RACE  ----------------------------------------
ORF2     ----------------------------------------
```

**Supplementary Figure 2. Sequence alignment of MALAT1 transcripts**

ClustalW alignment of the partial human MALAT1 transcript from the 5’ end, the MALAT1 decay intermediate identified previously by Mukherjee et al. (15), and the MALAT1 ORF2 sequence identified in this study. Nucleotides that are identical or conserved among the aligned sequences are highlighted according to the ClustalW colour scheme.

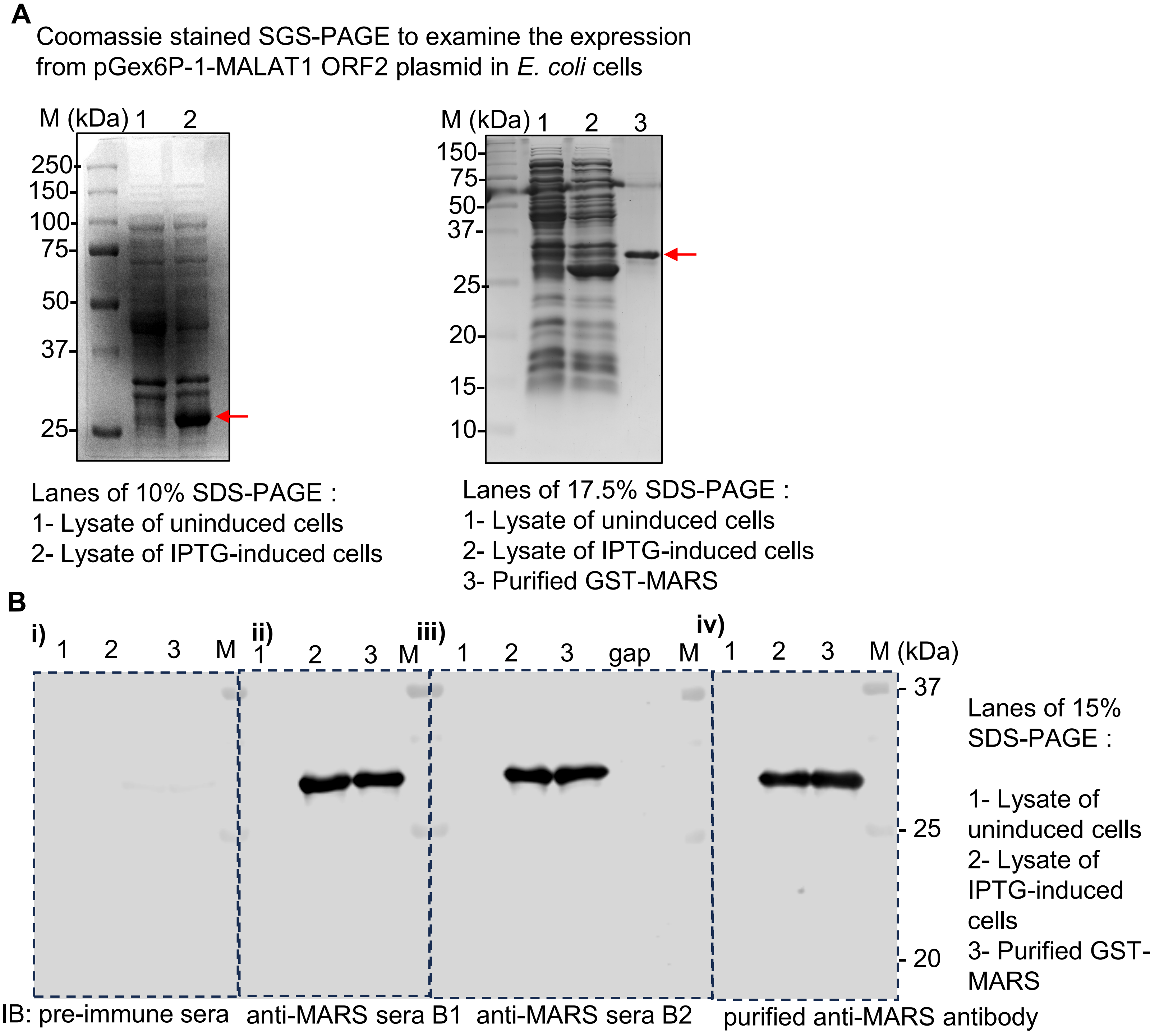


**Supplementary Figure 3. Validation and specificity of the anti-MARS antibody**

A) Coomassie-stained representative SDS-PAGE gels of different percentages showing expression of the recombinant GST-MARS chimeric protein in E. coli BL21DE3 cells. Lysates from uninduced and IPTG-induced cells expressing GST-MARS; and purified GST-MARS protein are analysed. The arrows indicate the expected protein bands corresponding to the recombinant fusion proteins. Molecular weight of the proteins is shown in the left. (B) Immunoblot analysis of the above protein samples with pre-immune sera, two antisera against MARS and purified anti-MARS antibody demonstrating the specificity of commercially raised anti-MARS antibody. Molecular weight of the proteins is shown in the right.

**A** ClustalW alignment of the sequences of human MALAT1 transcript (partial), mouse MALAT1 transcript (partial) and the human MALAT1 ORF2 identified in this study

```
                                   10        20        30        40        50        60        70
                          ....|....|....|....|....|....|....|....|....|....|....|....|....|....|
Mm                        CTCGCGACGGGTTCCGCGGTCCTCCCCGCCCTCCCTGGCGCGGCCCCCGCTTTCTGCGCCCAGTGACGCT
Hs                        ----------------------------------------------------------------------
ORF2                      ----------------------------------------------------------------------
Clustal Consensus
•••••
                                  220       230       240       250       260       270       280
                          ....|....|....|....|....|....|....|....|....|....|....|....|....|....|
Mm                        ATGGCCGCCAAGGTCGCCGTGCCCTCACCTGCAGCGACCATGGCCTTGCTGGGCTGAGACCGCAGCCTAA
Hs                        ------GTAAAGGACTGGGGCCCCGCAACTG--GCCTCTCCTGCCCTCTTAAGCGCAGCGC-CATTTTAG
ORF2                      ----------------------------------------------------------------------
Clustal Consensus
•••••
                                 2110      2120      2130      2140      2150      2160      2170
                          ....|....|....|....|....|....|....|....|....|....|....|....|....|....|
Mm                        CTTAATTTTGCTTGATGACAAAAGGACTTTTGATAACAGTTTCAAGATTGTCAGCATTTTGCATTGGACT
Hs                        TTAAACCGAAGGTGAT--TAAAAGACCTTGAAATCCATGACGCAGGGAGAATTGCGTCAT---TTAAAGC
ORF2                      ----------------------------------------------------------------------
Clustal Consensus

                                 2180      2190      2200      2210      2220      2230      2240
                          ....|....|....|....|....|....|....|....|....|....|....|....|....|....|
Mm                        TGAGCTGAGGTGCTTTTAAAATCCTAACGACTAGCATTGGCAGCTGACCCAGGTCTACACAGA--AGTGC
Hs                        CTAGTTAACGCATTTACTAAACGCAGACGA---AAATGGAAAGATTAATTGGGAGTGGTAGGATGAAACA
ORF2                      -----------------------------------ATGGAAAGATTAATTGGGAGTGGTAGGATGAAACA
Clustal Consensus                                            ** *  ** * *    **  *     **  *

                                 2250      2260      2270      2280      2290      2300      2310
                          ....|....|....|....|....|....|....|....|....|....|....|....|....|....|
Mm                        ATTCAGTGAACTAGGAAGACAGGAGCGGCAGACAGGAGTCCCGAAGC-CAGTTTGGTGAAGCTAGGAAGG
Hs                        ATTTGGAGAAGATAGAAGTTTGAAGTGGAAAACTGGAAGACAGAAGTACGGGAAGGCGAAG---AAAAGA
ORF2                      ATTTGGAGAAGATAGAAGTTTGAAGTGGAAAACTGGAAGACAGAAGTACGGGAAGGCGAAG---AAAAGA
Clustal Consensus         ***  * ***    ****   * ** ** * ** ***   * ****  * *   ** ****     ***

                                 2320      2330      2340      2350      2360      2370      2380
                          ....|....|....|....|....|....|....|....|....|....|....|....|....|....|
Mm                        ACTGAGGAGCCAGCAGCAGCAGTGCATGGTGAAGATAGCCCAGGAAAGAGTGCGGTTCGGTGGAGGAAGC
Hs                        ATAGAGAAGATAGGGAAATTAGAAGATAA-AAACATACTTTTAGAAGAAAAAAGATAAATTTAA---ACC
ORF2                      ATAGAGAAGATAGGGAAATTAGAAGATAA-AAACATACTTTTAGAAGAAAAAAGATAA------------
Clustal Consensus         *  *** **  **    *  **   **    ** ***      ***  *    * *

                                 2390      2400      2410      2420      2430      2440      2450
                          ....|....|....|....|....|....|....|....|....|....|....|....|....|....|
Mm                        TAGGAAGAAGGAGCCATACGGATGTGGTGGTGAAGCTGGGAAAGGGTTCCAGGATGGTGGAGCGAGAGCG
Hs                        TGAAAAGTAGGAAGCAGAAGAA---AAAAGACAAGCTAGGAAA------CAAAAAGCTAAGGGCAAAATG
ORF2                      ----------------------------------------------------------------------
Clustal Consensus
•••••
```

**B**

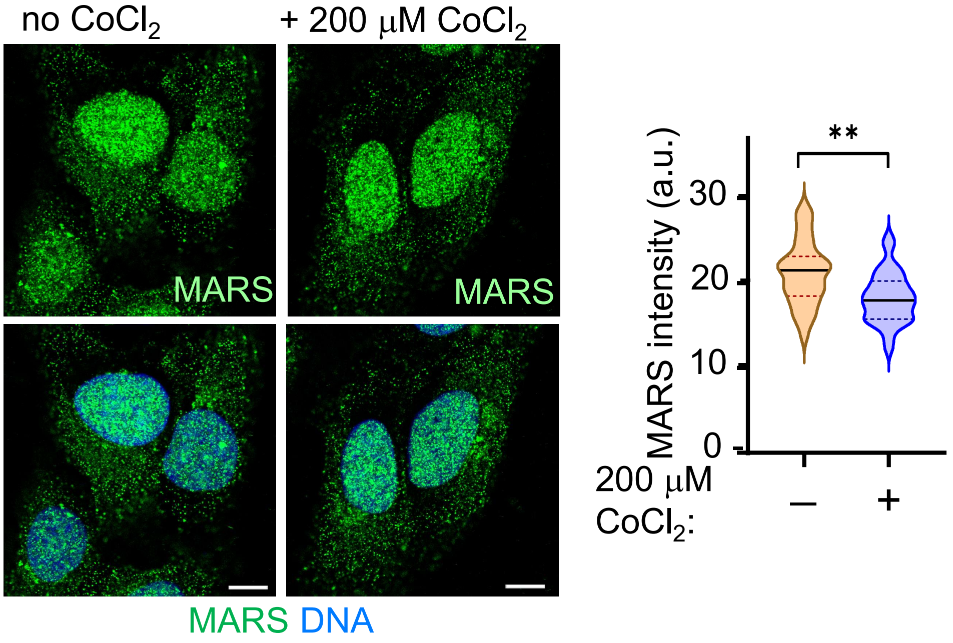

**Supplementary Figure 4. MARS is unique to human MALAT1**

(A) ClustalW alignment of partial human MALAT1, partial mouse MALAT1, and the human MALAT1 ORF2 sequence identified in this study. Conserved nucleotide positions are indicated above the alignment, and the consensus sequence is shown below. The alignment indicates that the ORF2 sequence identified in this study is specific to human MALAT1 and is not conserved in the corresponding mouse MALAT1 region. B) Immunofluorescence detection of MARS in U2OS cells under control conditions and following treatment with 200 μM $CoCl_2$ for 24 h resulted in a significant decrease in MARS fluorescence intensity compared with the untreated condition . Quantification of cellular MARS fluorescence intensity after background correction from 30 individual cells is shown on the right. MARS is shown in green and nuclear DNA in blue. The scale bar corresponds to 10 μm. Statistical significance is indicated by asterisks (*P < 0.05; **P < 0.01; ***P < 0.001), where ns represents non-significant.

### Gene Ontology analysis of MARS interactome

**A**

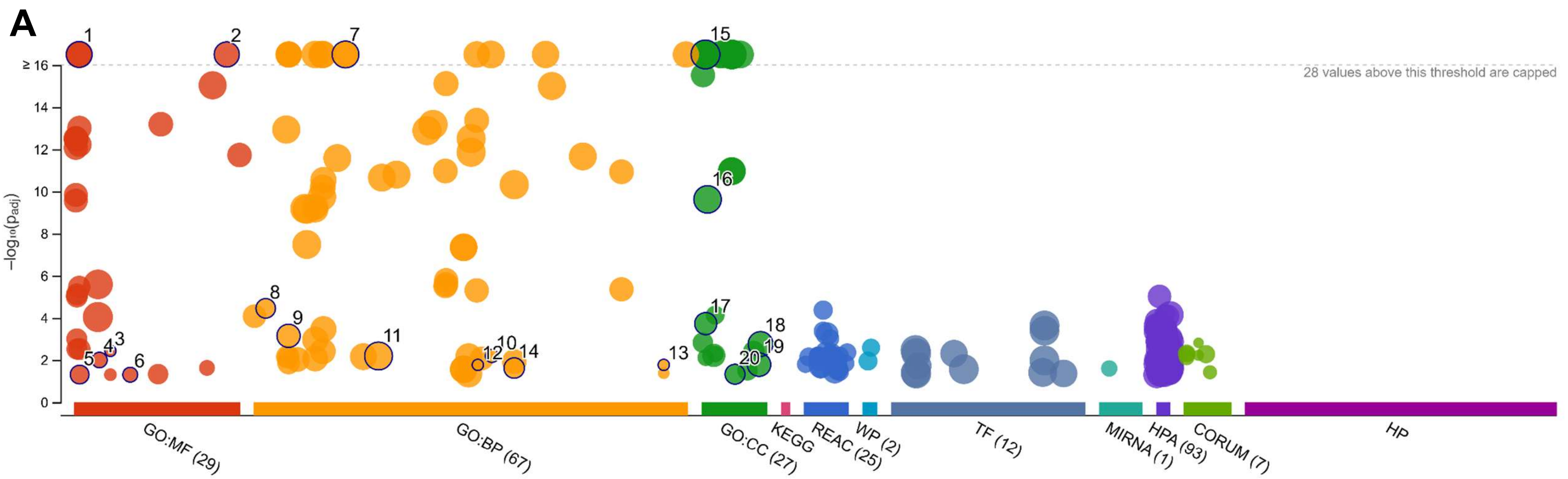


| ID | Source | Term ID | Term Name | $p_{adj}$ (query_1) |
|---|---|---|---|---|
| 1 | GO:MF | GO:0003677 | DNA binding | $1.047\times10^{-21}$ |
| 2 | GO:MF | GO:0140110 | transcription regulator activity | $1.693\times10^{-19}$ |
| 3 | GO:MF | GO:0010856 | adenylate cyclase activator activity | $3.698\times10^{-3}$ |
| 4 | GO:MF | GO:0008143 | poly(A) binding | $1.004\times10^{-2}$ |
| 5 | GO:MF | GO:0003730 | mRNA 3'-UTR binding | $4.916\times10^{-2}$ |
| 6 | GO:MF | GO:0019211 | phosphatase activator activity | $4.988\times10^{-2}$ |
| 7 | GO:BP | GO:0019219 | regulation of nucleobase-containing compound m... | $7.329\times10^{-32}$ |
| 8 | GO:BP | GO:0002181 | cytoplasmic translation | $3.541\times10^{-5}$ |
| 9 | GO:BP | GO:0006325 | chromatin organization | $7.116\times10^{-4}$ |
| 10 | GO:BP | GO:0060316 | positive regulation of ryanodine-sensitive calcium-... | $6.440\times10^{-3}$ |
| 11 | GO:BP | GO:0032502 | developmental process | $6.458\times10^{-3}$ |
| 12 | GO:BP | GO:0051343 | positive regulation of cyclic-nucleotide phosphodi... | $1.708\times10^{-2}$ |
| 13 | GO:BP | GO:1905913 | negative regulation of calcium ion export across pl... | $1.708\times10^{-2}$ |
| 14 | GO:BP | GO:0065004 | protein-DNA complex assembly | $2.409\times10^{-2}$ |
| 15 | GO:CC | GO:0005634 | nucleus | $5.567\times10^{-90}$ |
| 16 | GO:CC | GO:0005829 | cytosol | $2.388\times10^{-10}$ |
| 17 | GO:CC | GO:0005667 | transcription regulator complex | $1.879\times10^{-4}$ |
| 18 | GO:CC | GO:1902494 | catalytic complex | $1.737\times10^{-3}$ |
| 19 | GO:CC | GO:0140535 | intracellular protein-containing complex | $1.663\times10^{-2}$ |
| 20 | GO:CC | GO:0044391 | ribosomal subunit | $4.733\times10^{-2}$ |

**B**

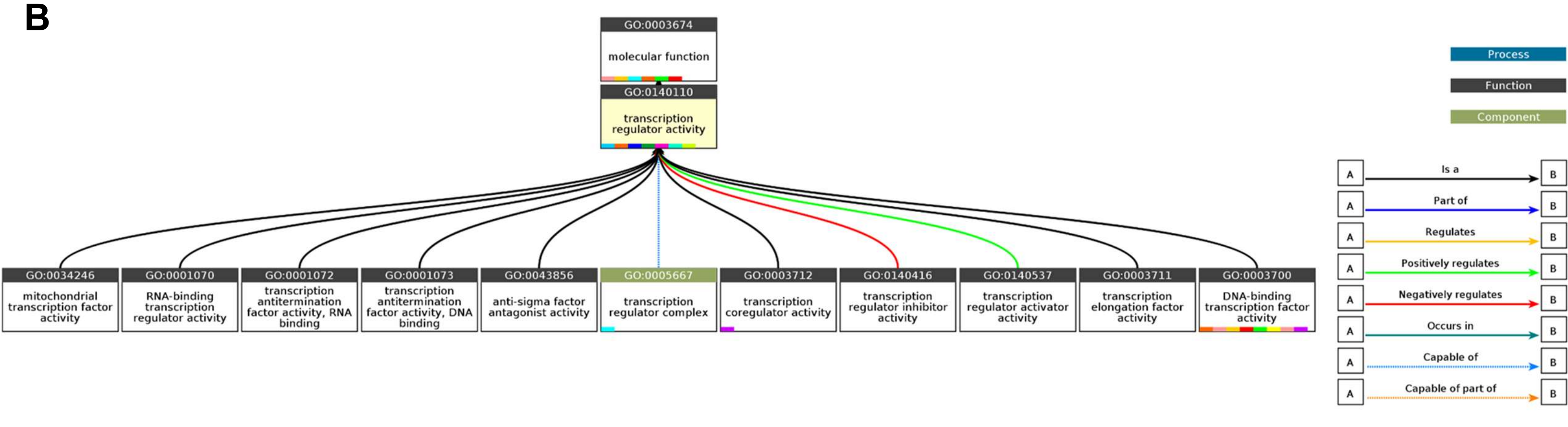


**Click the graph to show a high resolution version.**

(a) Cscore$^{GO}$ is the confidence score of predicted GO terms. Cscore$^{GO}$ values range in between [0-1]; where a higher value indicates a better confidence in predicting the function using the template.

(b) The graph shows the predicted terms within the Gene Ontology hierachy for Cellular Component. Confidently predicted terms are color coded by Cscore$^{GO}$:

[0.4,0.5) [0.5,0.6) [0.6,0.7) [0.7,0.8) [0.8,0.9) [0.9,1.0]

**Supplementary Figure 5. Identification of MARS interactome**

(A) Gene Ontology (GO) enrichment analysis of proteins identified in the MARS interactome. The bubble plot summarizes the enriched GO terms across the major biological process, cellular component, and molecular function categories. The position and size of the bubbles represent the corresponding enrichment values and number of proteins associated with each term, respectively, while the colour scale indicates the significance of enrichment. The table below lists the representative enriched GO terms together with their corresponding GO identifiers, functional descriptions, and enrichment values. (B) Functional interaction network generated from the proteins identified in the MARS interactome. The network groups the interacting proteins into functionally related clusters and highlights the major cellular processes and pathways associated with the MARS-interacting protein set. The different branches represent the functional categories identified from the interaction analysis, providing an overview of the biological functions represented within the MARS interactome.

Molecular docking study of MARS and its interacting protein Nrf2

Nrf2 vs MARS interaction

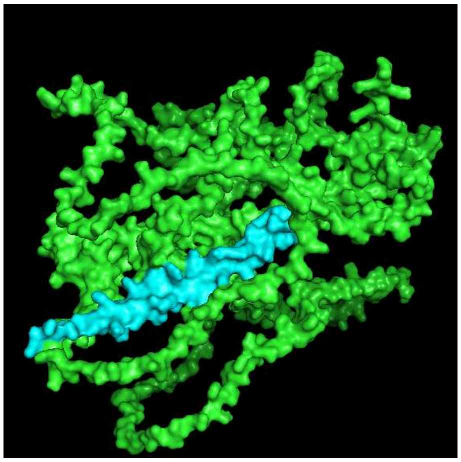

| | |
|---|---|
| HADDOCK score | -107.8+/- 4.0 |
| Cluster size | 14 |
| RMSD from the overall lowest-energy structure | 1.2 +/- 0.7 |
| Van der Waals energy | -37.7 +/- 11.8 |
| Electrostatic energy | -378.1 +/- 36.5 |
| Desolvation energy | -8.5 +/- 4.4 |
| Restraints violation energy | 585.3 +/- 97.5 |
| Buried Surface Area | 1781.0 +/- 32.6 |
| Z-Score | -1.8 |

**Supplementary Figure 6. Molecular docking analysis predicts potential interactions of MARS with Nrf2**

(A) Molecular docking was performed to examine the possible interaction between MARS and Nrf2. The predicted MARS-Nrf2 complex is generated as a space filled molecular surface representation in PyMOL, with Nrf2 and MARS indicated in green and cyan, respectively. The docking solution was evaluated using the HADDOCK scoring parameters, and the lowest-energy cluster was selected for visualization. The corresponding cluster size, RMSD from the lowest-energy structure, van der Waals energy, electrostatic energy, desolvation energy, restraint violation energy, buried surface area, and Z-score are shown alongside the docked structure. The docking analysis predicts a direct physical association between MARS and Nrf2 and provides a structural model for the potential interaction between the two proteins. The analysis was performed using the HADDOCK web server.

MARS can bind to the Antioxidant Response Element (ARE) of DNA

Docking of MARS and ARE dsDNA [TGCTGA(G/C)TCAGCA]

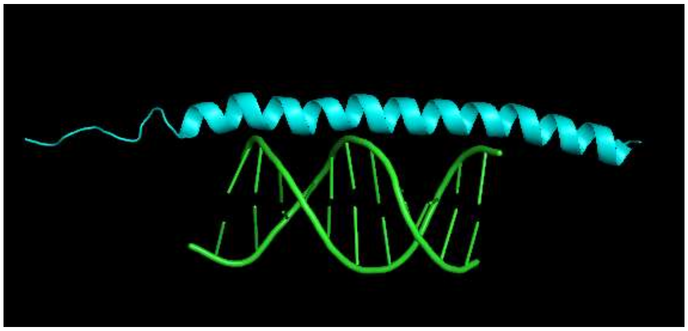

| | |
|---|---|
| HADDOCK score | -83.3 +/- 9.5 |
| Cluster size | 19 |
| RMSD from the overall lowest-energy structure | 1.2 +/- 0.8 |
| Van der Waals energy | -57.2 +/- 6.7 |
| Electrostatic energy | -378.1 +/- 36.5 |
| Desolvation energy | -9.0 +/- 4.4 |
| Restraints violation energy | 585.3 +/- 97.5 |
| Buried Surface Area | 1781.0 +/- 32.6 |
| Z-Score | -1.8 |

Docking of MARS and HRE dsDNA [A/GCGTG]

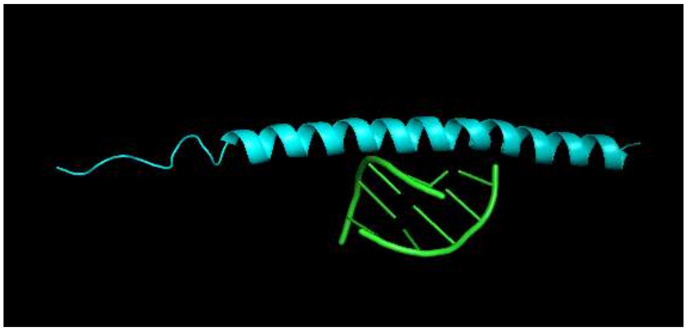

| | |
|---|---|
| HADDOCK score | 4.9 +/- 9.0 |
| Cluster size | 4 |
| RMSD from the overall lowest-energy structure | 7.2 +/- 0.4 |
| Van der Waals energy | -27.7 +/- 7.5 |
| Electrostatic energy | -198.8 +/- 33.7 |
| Desolvation energy | -0.3 +/- 1.6 |
| Restraints violation energy | 727.2 +/- 46.1 |
| Buried Surface Area | 974.9 +/- 75.3 |
| Z-Score | 1.3 |

https://rascar.science.uu.nl/haddock2.4/

**Supplementary Figure 7. Molecular docking analysis predicts potential interactions of MARS with antioxidant response element (ARE)-containing DNA**

Molecular docking analysis showing the predicted interaction of the MALAT1 ORF2-encoded peptide MARS with the ARE of double stranded (ds) DNA. ARE dsDNA sequence used for docking is indicated above the structure. The predicted MARS-ARE dsDNA complex is shown as a molecular surface representation, with MARS and DNA highlighted separately. The corresponding HADDOCK score, cluster size, RMSD from the lowest-energy structure, van der Waals energy, electrostatic energy, desolvation energy, restraint violation energy, buried surface area and Z-score are indicated below the structure. Molecular docking of MARS with the HRE dsDNA sequence, used as a control for comparison with the ARE-containing DNA. The predicted MARS-HRE dsDNA complex and the corresponding HADDOCK parameters are shown. The docking analysis suggests a more favourable interaction of MARS with the ARE sequence compared with the HRE sequence, supporting the possibility that MARS may associate preferentially with the ARE-containing DNA.

| Oligo nomenclature | Oligo sequences |
|---|---|
| pEGFPN1 mut_FP | CACCTAATTGTTAGCAAGGGCGAGGAGCTG |
| pEGFPN1 mut_RP | GCTAACAATTAGGTGGCGACCGGTGGATC |
| MALAT1 ORF2 ATG mut FP | GCTTTCCATTGCGATTTGCCTTGTGAGCACT |
| MALAT1 ORF2 ATG mut RP | ATCGCAATGGAAAGCTTGAGCTCGAGATCTGA |
| MALAT1 ORF7 ATG mut FP | GCTTAAAATTGAAAGATTAATTGGGAGTGGTAGGA |
| MALAT1 ORF7 ATG mut RP | CTTTCAATTTTAAGCTTGAGCTCGAGATCTGAG |
| L10A pLVX FP | TAAGAATTCACCATGTACCCATACGATGTTCC |
| L10A pLVX RP | ATTGGATCCTTAATATAGGCGCTGGGG |
| CAT ARE 1 FP | TGCTTACTCCTGACACCACG |
| CAT ARE 1 RP | AGTGCTGAGAGAAGGGAAGG |
| CAT ARE 2 FP | AGTCTTGCTCTGTCACCCG |
| CAT ARE 2 RP | CTTGCCTGTAGTCCCAGCT |
| CAT ARE 3 FP | CCACGTCCTTTGCTGTTGTA |
| CAT ARE 3 RP | TCAGCCATAACCACTAACCCA |
| SOD2 ARE 1 FP | AGTGTTGGGATTACAGGCGT |
| SOD2 ARE 1 RP | TGACCCCTGAATGATGCAGT |
| SOD2 ARE 2 FP | ACCACAGGCAGGAAGAAAGA |
| SOD2 ARE 2 RP | TCGTTGTTGCTAAATATCCTCCT |
| CAT control_FP | CCATGAGGCAGCTTTGAAGA |
| CAT control RP | TCCAGTAACAGATCCAACCACA |
| SOD control FP | CGGGAAAGATATGGTGCAGC |
| SOD control RP | AGCCAAAGAAGAAGGTGGGA |
| MALAT1 sORF1 FP | ACGAAGCTTTCCATGGCGATTTGCC |
| MALAT1 sORF1 RP | CGTGGATCCGAAACGTGAAAACCCACT |
| MALAT1 sORF2 FP | ACGAAGCTTAAAATGGAAAGATTAATTGGG |
| MALAT1 sORF2 RP | CGTGGATCCTCTTTTTTCTTCTAAAAGTATG |
| RP1 ORF1 FP | ACGAAGCTTTCCATGTTCCCCACGT |
| RP1 ORF1 RP | CGTGGATCCGAACTTGAGGCTGAA |
| RP1 ORF2 FP | ACGAAGCTTCTGATGATCCAGCTGCTGC |
| RP1 ORF2 RP | ACGGGTACCCTGGGGGTCCCCAGCT |
| GAS5 ORF1 FP | ACGAAGCTTAGGATGAGAATAGC |
| GAS5 ORF1 RP | CGTGGATCCACTTCCAGCTTTCT |
| GAS5 ORF2 FP | ACGAAGCTTAGAATGGAGTGCAGCG |
| GAS5 ORF2 RP | CGTGGATCCCTTGAGGTCAAGAGTT |
| MALAT1 control G4 | AGGGAAGGGAGGGGGTGCCTGTGGGG |
| MALAT1 RACE product FP inf | TCTCGAGCTCAAGCTTGCTTAGCGTTAAGTTTTTAACGT |
| MALAT1 RACE product RP inf | TTTCCATTTTAAGCTCGTCTGCGTTTAGTAAATGCGT |
| MALAT1 5'-end FP inf | TCTCGAGCTCAAGCTGTAAAGGACTGGGGCCCCG |
| MALAT1 5'-end RP new1 inf | AAGTGGCAAAATGGCGGACTTTCTCCCCC |
| MALAT1 5'-end FP new1 inf | AGTCCGCCATTTTGCCACTTCTCAACCGTCC |
| Seq primer pEGFPN1 | CATGGCGATTTGCCTTGTGAGCAC |
| FP_G42 at G41 | GCTGAGGGGGCAGGCGGAGCTTGAGGACACTGGCAGCC AACGGCC |
| RP_G42 at G41 | GCCTGCCCCCTCAGCAGAAGCCCCTCACGGAAATTTTTCTA CCGTTTTTC |
| FP_G41 at G42 | GCGGCAACTGGGGGGCCGCAGATCAGAGTGGGCAACCG CAGATAAGTTTTTTTCTC |
| RP_G41 at G42 | CCCCCCAGTTGCCGCCAGCCCCCCACGGCCCGGCGCTGGC TGCCTCAATG |

| | |
|---|---|
| FP_NRAS CDS pEGFPN1 mut | ATTCTGCAGTCGACGGTACCATGACTGAGTACAAACTGGT |
| RP_NRAS RNA pEGFPN1 mut | GGCGACCGGTGGATCCCATCACCACACATGGCAATCCC |
| FP_NRAS RNA pEGFPN1 mut | ATTCTGCAGTCGACGGTACCGAAACGTCCCGTGTGGGAG G |
| FP_NRAS G4 mut in pEGFPN1 mut | AAAAAAACAAATCTAAATGCGGCCTGCCGCATGACTCG |
| RP_NRAS G4 mut in pEGFPN1 mut | TAGATTTGTTTTTTTTACACGGGACGTTTCGGTACC |
| FP_NRAS to MALAT1 rG41 | GCGGCAACTGGGGGGCCGCAGATCAGAGTGGGCGCGGC CTGCCGCATGACTCG |
| RP_NRAS to MALAT1 rG41 | CCCCCCAGTTGCCGCCAGCCCCCCACGGCCCGCACGGGAC GTTTCGGTACCG |
| FP_NRAS to MALAT1 rG42 | GCTGAGGGGGCAGGCGGAGCTTGAGGAGCGGCCTGCCG CATGACTCG |
| RP_NRAS to MALAT1 rG42 | GCCTGCCCCCTCAGCAGAAGCCCCTCACGGGACGTTTCGG TACCG |
| FP_HA-RPL10A in pCDNA4TO | TTAAACTTAAGCTTGGTACCATGTACCCATACGATGTTCCA GATTACGCTAGCAGCAAAGTCTCTCGCGACACCC |
| RP_RPL10A in pcDNA4TO | CTGGACTAGTGGATCCTTAATATAGGCGCTGGGGCTTG |
| FP_HCV IRES in sORF2 | CGTCAGATCCGCTAGCTCTAGAACTAGTGGATCCCCCGGG |
| RP_HCV IRES in sORF2 | TTTCCATTTTAAGCTTGACGTCCTGTGGGCGGCG |
| FP rG4 1 mut | ACAGCAACTGAAAAGCCGCAGATCAGAGTGAGCCACTGG CAGCCAACGGC |
| RP rG4 1 mut | CTTTTCAGTTGCTGTCAGCTTTTCACGGCTCGCACGGAAAT TTTTCTACCGTTT |
| FP rG4 2 mut | GCTGAAAAAGCAGACAGAGCTTGAAGAAACCGCAGATAA GTTTTTTTCTCT |
| RP rG4 2 mut | GTCTGCTTTTTCAGCAGAAGCTTCTGCGCTGGCTGCCTCAA T |
| FP_rG42 in pRLTK inf | GCTGAGGGGGCAGGCGGAGCTTGAGGTCTAGAGCGGCC GCTTCG |
| RP_rG42 in pRLTK inf | GCCTGCCCCCTCAGCAGAAGCCCCGCTAGCCTATAGTGAG TCGT |
| FP_non rG4 in pRLTK inf | AGGTGATCGAATTCCGGTGATGCGAGTCTAGAGCGGCCG CTTCG |
| RP_non rG4 in pRLTK inf | GGAATTCGATCACCTTCCGCCGCCGCTAGCCTATAGTGAG TCGT |
| FP_MALAT1 full | AGGTTTCCCAGAGTCCTTGG |
| RP_MALAT1 full | GCTTTCCTCCAAACCCCAAG |
| FP_MALAT1 decay | GCAGGGAGAATTGCGTCATT |
| RP_MALAT1 decay | TTCTTCGCCTTCCCGTACTT |
| FP_NFYC ORF2 clone pEGFP inf | TGCTGATGGTAACCTTGAATTTTACAAAGCGGGCCCGGGA TCCACC |
| RP_NFYC ORF2 clone pEGFP inf | AGGTTACCATCAGCAGAGGCAGTCATGGTACCGTCGACTG CAGAATTCG |
| FP_NFYC ORF1 clone pEGFP | TTAGGTACCATGAATGAACAAAACCTA |
| RP_NFYC ORF1 clone pEGFP | AATGGATCCTTAGATCTTTTTACAGATG |
| FP_MALAT1 rG4 swap mut1 | TCTGCAACTGTTTTGCCGCAGATCAGAGTTTGCAACCGCA GATAAGTTTTTTTCTCT |
| RP_MALAT1 rG4 swap mut1 | CAAAACAGTTGCAGACAGAAAAAAACGGAACGGCGCTGG CTGCCTCAAT |

| | |
|---|---|
| FP_MALAT1 rG4 swap mut2 | GCTGATTTTGCATGCGTAGCTTGATGACACTGGCAGCCAACGGC |
| RP_MALAT1 rG4 swap mut2 | GCATGCAAAATCAGCAGAAGCAACTCACGGAAATTTTTCTACCGTTTT |
| FP_NRAS RNA pEGFPN1 mut mod | ATTCTGCAGTCGACGGTACCGAAACGTCCCGTGTGGGAGGGGCGGGTCTGGGTGCGGCCTGCCGCATG |
| FP_rG42 in pRLTK inf | GCTGAGGGGGCAGGCGGAGCTTGAGGTCTAGAGCGGCCGCTTCG |
| RP_rG42 in pRLTK inf | GCCTGCCCCCTCAGCAGAAGCCCCGCTAGCCTATAGTGAGTCGT |
| FP_non rG4 in pRLTK inf | AGGTGATCGAATTCCGGTGATGCGAGTCTAGAGCGGCCGCTTCG |
| RP_non rG4 in pRLTK inf | GGAATTCGATCACCTTCCGCCGCCGCTAGCCTATAGTGAGTCGT |
| CAT qPCR_FP | AGAAGACTCCCATCGCAGTT |
| CAT qPCR_RP | TTTCACTGCAAACCCACGAG |
| SOD2 qPCR_FP | CTGGAACCTCACATCAACGC |
| SOD2 qPCR_RP | CCTGGTACTTCTCCTCGGTG |
| PRDX2 qPCR_FP | ACTCTCAGTTCACCCACCTG |
| PRDX2 qPCR_RP | ACGCCGTAATCCTCAGACAA |
| GLRX1 qPCR_FP | GTGTTCATCAAGCCCACCTG |
| GLRX1 qPCR_RP | GTTAGTGTGGTTGGTGGCTG |
| GPX4 qPCR_FP | GCCAGGGAGTAACGAAGAGA |
| GPX4 qPCR_RP | CTTCATCCACTTCCACAGCG |
| GCLC qPCR_FP | CTCCAGGTGACATTCCAAGC |
| GCLC qPCR_RP | CGGTAAAAGGGAGATGCAGC |

**Supplementary Table-1**: Oligos used in this study
